# *HLF* Expression Defines the Human Hematopoietic Stem Cell State

**DOI:** 10.1101/2020.06.29.177709

**Authors:** Bernhard Lehnertz, Jalila Chagraoui, Tara MacRae, Elisa Tomellini, Sophie Corneau, Nadine Mayotte, Isabel Boivin, Guy Sauvageau

## Abstract

Hematopoietic stem cells (HSCs) sustain blood cell homeostasis throughout life and are able to regenerate all blood lineages following transplantation.

Despite this clear functional definition, highly enriched isolation of human HSCs can currently only be achieved through combinatorial assessment of multiple surface antigens. While a number of transgenic HSC reporter mouse strains have been described, no analogous approach to prospectively isolate human HSCs has been reported.

To identify genes with the most selective expression in human HSCs, we profiled population- and single-cell transcriptomes of fresh and *ex vivo* cultured cord blood derived HSPCs as well as peripheral blood, adult bone marrow and fetal liver. Based on these analyses, we propose the master transcription factor *HLF* (*Hepatic Leukemia Factor*) as one of the most specific HSC marker genes.

To directly track its expression in human hematopoietic cells, we developed a genomic *HLF* reporter strategy, capable of selectively labeling the most immature blood cells on the basis of a single engineered parameter.

Most importantly, *HLF*-expressing cells comprise all of the stem cell activity in culture and *in vivo* during serial transplantation.

Taken together, these results experimentally establish *HLF* as a defining gene of the human hematopoietic stem cell state and outline a new approach to continuously mark these cells with high fidelity.

**Key Points:**

- In the human blood system, *HLF* expression is specific to stem cell populations in primary anatomical sites and during *ex vivo* expansion.
- CRISPR/rAAV6-mediated integration of a genomic *HLF*-reporter allows selective and stable genetic labeling of human HSCs *ex vivo* and *in vivo*.

## Introduction

The existence of rare, serially transplantable and multipotent hematopoietic stem cells (HSCs) was first demonstrated nearly sixty years ago in mice^1–3^. Since then, HSCs have been studied extensively not only due to their unique biology but because of their paramount regenerative potential that is now widely exploited in the clinic^4^.

Despite their high clinical relevance, the molecular identity of human HSCs remains poorly defined and their purification invariably requires profiling of complex stem-cell associated surface marker combinations^5^. Since many of these surface markers are mediators of cellular homing or signaling, their expression is often regulated in response to changing physiological conditions, significantly impacting utility during certain experimental procedures, most notably *ex vivo* culture^6,7^.

In mice, several HSC-enriched genes encoding intracellular proteins have been identified and, using transgenic reporter strains, were demonstrated to label repopulating cells with high accuracies^8–14^. Collectively, these studies suggest that a number of intracellular proteins, particularly transcriptional and chromatin regulators, are more specifically expressed than most, if not all, currently used surface HSC markers.

With recent advancements in targeted gene editing using CRISPR and recombinant adeno-associated viruses (rAAV)^15^ and in functional expansion of human HSCs in culture^16–20^, the use of genetic reporter alleles in these cells has become conceivable. Moreover, improved characterization of developmental gene expression networks using single cell transcriptomics has set the stage for population marker gene identification with unmatched resolution. Through combination of these key advancements, we set out to identify the most selectively expressed candidate genes in human HSCs and engineer a genomic reporter allowing prospective identification of *bona fide* human HSCs in culture and *in vivo*.

## Methods

### Analyses of bulk transcriptomes

EPCR and ITGA3 specific datasets were previously reported. In brief, differentially expressed genes (DEGs) from the ITGA3 dataset were determined exactly as in^21^ using the Kallisto/Sleuth pipeline and the full GRCh38 v92 annotation (including non-coding genes). The EPCR dataset^7^ was re-analyzed in the same fashion for consistency. Expression weighted fold-change (beta-value) and p-value (sleuth) cut-off values were designated based on ITGA3 (beta ± 0.844; p = 0.0071) and EPCR (beta ± 1.412; p = 3.32e-7) in the respective datasets. Intersection of positive and negative DEGs yielded 17 and 7 genes respectively. Analyses and heatmaps were generated in R.

### Analyses of single cell transcriptomes from fresh and UM171 expanded cord blood

CD34^+^ cord blood (CB) cells, either freshly thawed or culture-expanded with UM171 (35nM) were single-cell sequenced on a Chromium Single-Cell Controller (10X Genomics) using the Single Cell 3’ Reagent Kit version 2 according to manufacturer’s instructions. Target cell numbers were 6,000 per condition. scRNAseq libraries were sequenced on an Illumina NovaSeq device using a S2 (PE 28×91) setup or on an Illumina HiSEQ 4000 using 26×98 cycles. A standard Cellranger v3.0.1 pipeline was used for read mapping (GRCh38 annotation) and demultiplexing. Subsequent analyses were done in Seurat (v3) based on Cellranger prefiltered barcode/feature matrices and included (i) exclusion of cells with less genes or UMIs than the respective medians minus 2 standard deviations (ii) exclusion of cells with more genes or UMIs than the respective medians plus 2 standard deviations (multiplets), (iii) exclusion of cells with more than 7% mitochondrial gene expression (apoptotic cells). Expression counts were normalized using the SCTransform wrapper in Seurat including regression on cell cycle scores and mitochondrial gene content. Seurat integration was performed using the top 241 integration anchors (250 minus sex specific genes) in the first 30 dimensions, followed by PCA dimensional reduction, FindNeighbors and FindClusters (resolution = 0.5) in the first 15 integrated dimensions. SPRING embedding was calculated on the integrated expression matrix using the SPRING webtool (https://kleintools.hms.harvard.edu/tools/spring.html). For visualization, data imputation was calculated on SCT transformed data of all genes using the MAGIC wrapper (t = 1) in Seurat.

### CD34^+^ enriched cord blood cell culture

Culture of isolated human CB-derived CD34+ cells was performed in HSPC expansion media comprised of Stem Span serum-free media (StemCell Technologies #09855) supplemented with Glutamax (Invitrogen #35050061), low density lipoprotein (StemCell Technologies #02698), 100 ng/ ml SCF (Shenandoah Biotechnology #100-04), 100 ng/ml Flt3L (Shenandoah Biotechnology #100-21), 50 ng/ml TPO (R&D system #288-TP) and 35 nM UM171 (StemCell Technologies #72914).

### Nucleofection and transduction of CD34^+^ cells

Nucleofection of was carried out with 3ug Cas9 and 8ug of sgRNA and 10e6 cells per 100 ul, using the DZ100 program. 4 days before nucleofection, CD34+ cells were thawed and plated at 1.5*10e5 cells/ml in HSPC expansion media containing UM171 (35 nM). At day 3, cells had typically expanded 2-3fold, and were stained with anti-CD34-BV421 (BD #562577, 1:50) and anti-CD201-APC (Biolegend #351906, 1:100) and sorted based on CD201 expression. The sorted cells were plated back in culture at 1.5*10e5 cells/ml. 24h later, cells were harvested, washed in PBS and taken up in 1M nucleofection buffer containing 11ug the pre-assembled Cas9 sgRNA RNP complex and were indicated 20 fmol of siRNA against TP53 (ThermoFisher #4390825, siRNA id s605). After nucleofection, the cells were immediately plated at 2-4*10e5 cell per ml in HSPC media optionally containing 400 MOIs of reporter encoding rAAV6. Half media changes were done on days 5 and 6, analysis or transplantation was done on day 7.

### Assessment of homologous recombination using droplet digital PCR (ddPCR)

Primers were designed to amplify a 605bp region of the wildtype or 601bp region of the targeted allele using a set of 3 primers. A common external forward primer (ext_FW; 5’-CCACCTGCTTTCATCCAGC-3’) binds in intron 3 upstream of the left homology arm, while the reverse primer binds either in the right homology arm (3’_RV; 5’-GGTAAAGTGCTGATGTCAGAAAGG-3’) or in the IRES region of the reporter (ires_RV; 5’-TAACATATAGACAAACGCACACCG-3’). The 3’_RV primer binds 77bp downstream of the Cas9 cut site, thus amplifying most non-integrated alleles. A FAM labeled non-fluorescent quencher dual-labeled probe (HLF_common_FAM_probe; 5’-FAM-TCTGATCTCTGCTTCACTGAGCACGC-ZEN-Iowa-black-3’) binds the HLF locus within the left homology arm and as such will detect both the wildtype and targeted allele amplicons. A second probe labelled with HEX and non-fluorescent quencher (ires_HEX_probe; 5’-HEX-CAAGCGGCTTCGGCCAGTAACGTTAG-ZEN-Iowa-black-3’) binds in the IRES cassette and will detect only targeted allele amplicons. PCR efficiency and specificity were validated in standard PCR assays prior to ddPCR using synthetic gBlock DNA fragments consisting of synthetic wildtype or targeted alleles, combined with primer drop-out testing. Oligonucleotides and gBlocks were synthesized by IDT. To assess targeting of the HLF locus following electroporation and in xenograft recipients, approximately 30,000 cells were lysed in 25ul Quick Extract Buffer (Lucigen #QE09050), according to the manufacturer’s directions. Briefly, cells were pelleted and resuspended in lysis buffer, then heated at 65° for 6min, vortexed and incubated for 2min at 95°. For cell line experiments with a larger cell numbers, the amount of lysis buffer was scaled up accordingly. 22ul ddPCR reactions contained 2.2ul of cell lysate, 11ul of 2X ddPCR Supermix for Probes (No dUTP, BioRad #1863024), 1.1ul (4U) FastDigest-HindIII (prediluted 1/3) (Thermo Fisher) and 11.1ul 20X primer-probe mix. Final concentration of primers and probes were 0.25uM and 0.9uM each, respectively. ddPCR reaction emulsion was created in a DG8 cartridge with DG8 gasket (BioRad #1863009) from 20ul PCR mix and 70ul droplet generation oil for probes (BioRad #1863052), using a QX200 Droplet Generator (BioRad #1864002). 40ul of PCR emulsion was pipetted into a 96-well QX200 compatible PCR plate (BioRad #12001925), which was covered with a Pierceable Foil Heat Seal and sealed in a PX1 PCR plate sealer (BioRad #1814000) before cycling in a C1000 deep-well thermocycler (BioRad #1851197). Thermocycler conditions were as follows: 95°C for 10min; 50 cycles of 30sec at 95°C, 2min at 58°C, 2min at 72°C, followed by 10 min at 98°C, with a ramp speed of 2°C / step throughout. ddPCR products were measured using a QX200 Droplet reader (BioRad #1864001). Amplitude and Cluster data was exported from QuantaSoft Software (BioRad #1864003) and analyzed in R using the ddPCR package.

### Data Availability

The accession numbers for the previously published datasets are as follows: EPCR population dataset (GSE77128), ITGA3 population dataset (GSE130974), the human bone marrow single cell dataset is available through the Human Cell Atlas consortium and was downloaded as count matrix through the HCAData portal in R (https://github.com/federicomarini/HCAData). The human fetal liver dataset was obtained as annotated count matrix from Muzlifah Haniffa or else is available at ArrayExpress with accession code E-MTAB-7407. Fresh and UM171-expanded CD34+ single-cell datasets have been deposited to GEO under accession GSE153370.

### Additional methods

All additional methods are provided in the supplemental Data.

## Results

### *Hepatic Leukemia Factor* (*HLF*) is a candidate marker gene for human HSC populations

We previously identified surface markers such as EPCR (CD201) and ITGA3 (CD49c) that best define long-term repopulating cells in optimized *ex vivo* CD34^+^ cord blood stem cell expansion conditions^7,21^.

Integrated transcriptome analysis of these enriched (CD34^+^/CD201^+^ and CD34^+^/CD45RA^low^/CD201^+^/CD90^+^/CD133^+^/ITGA3^+^) versus depleted LT-HSC populations yielded a set of genes (n=17) with strongly LT-HSC-associated expression (**Supplemental Fig. S1**). Based on expression dynamics between LT-HSCs and differentiated cells, *Hepatic Leukemia Factor* (*HLF*) ranked highest in this list and was therefore prioritized as an HSC marker candidate (**Fig. 1A**). Indeed, *HLF* was found not expressed in mature peripheral blood cells^22^ (**Fig. 1B**) and single-cell transcriptomes of freshly isolated and culture-expanded CD34^+^ cord blood cells confirmed its expression in the hematopoietic stem and progenitor cell (HSPC) cluster (**Fig. 1C, pink cluster and Fig. 1D and E**). Moreover, within this cluster, *HLF* expression continuously decreased as cells progressed towards lineage commitment, a pattern that aligns with latest models of gradual rather than stepwise HSC differentiation^23,24^.

**1).**
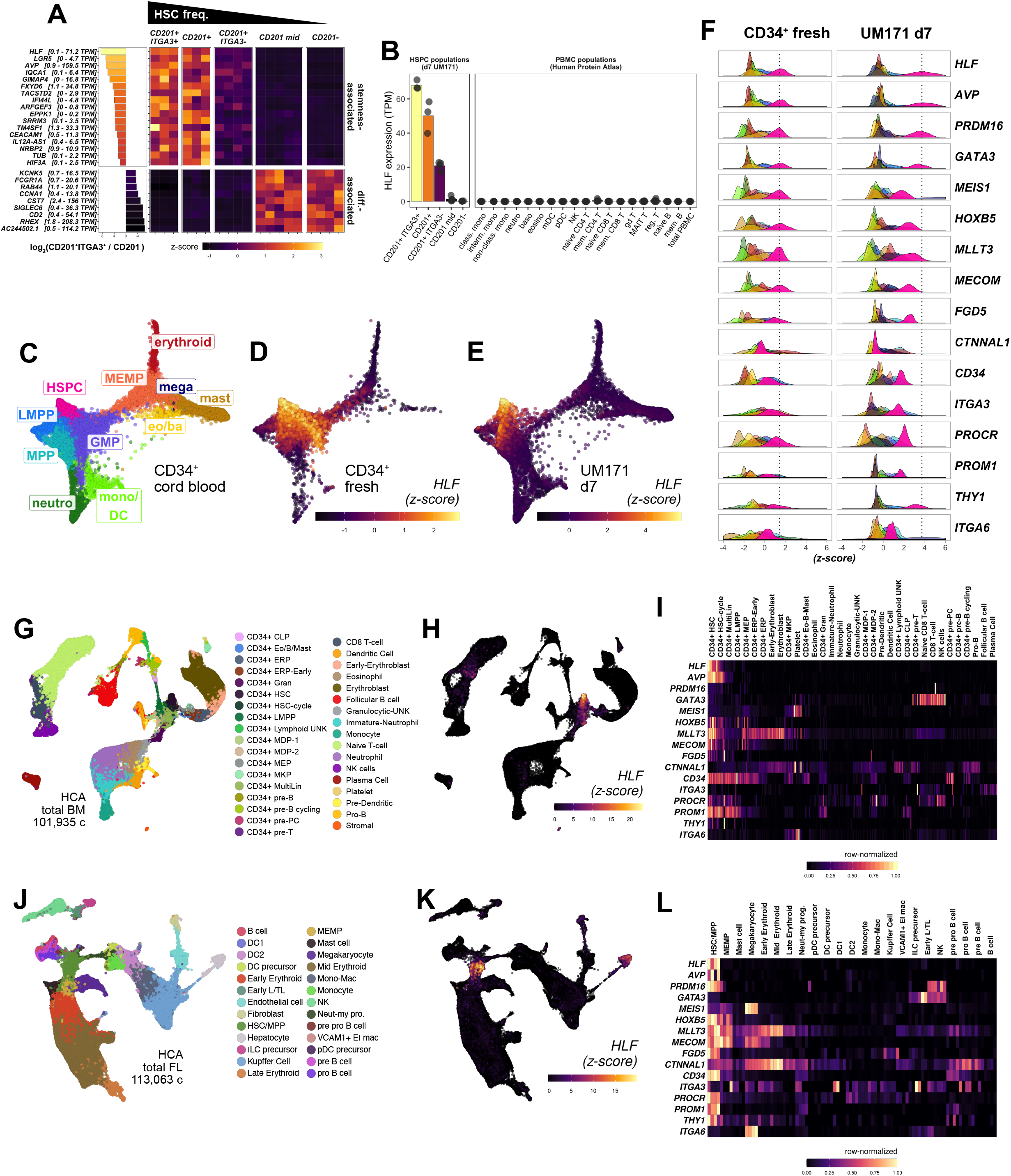
Specific *HLF* Expression in Enriched Human HSC Populations. ***A) HLF expression is enriched in cultured human HSC subsets*.** Differentially expressed genes from CD201 + ^7^ and ITGA3/CD201+ ^21^ HSC-enriched population transcriptomes were intersected to identify consistently up or downregulated genes. Ranking based on fold-change between the most enriched (ITGA3^+^/CD201^+^) and most depleted (CD201-) HSC populations is summarized by a waterfall plot (left, log2-transformed). Range of expression is provided for each gene in square brackets (TPM = transcripts per kilobase million). Each population includes biological replicates represented along the x-axis. ***B) HLF-expression is undetectable in blood leukocyte populations*.** Curated dataset from^22^, between 4 and 6 biological replicates per population. ***C) Single cell transcriptomic overview of cord blood cell populations*.** Fresh and UM171-expanded CD34^+^ (d7) (2 biological replicates for each condition, 15,921 cells total), were scRNA sequenced (10X Chromium), integrated and clustered using Seurat 3^47^). Ten cell clusters were identified: hematopoietic stem/progenitor cells (HSPC), lymphoid-primed multipotent progenitors (LMPP), multipotent progenitors (MPP), granulo-monocytic progenitor (GMP), megakaryocyte-erythroid-mast cell progenitors (MEMP), megakaryocytes (mega), eosinophil/basophils (eo/ba), mast cells (mast), erythroid lineage cells, neutrophils (neutro) and monocytic/dendritic cells (mono/dc). Dimensional reduction was calculated using SPRING ^48^. ***D-E) HSPC specific HLF expression in fresh CD34^+^ cells and after 7d ex vivo expansion in the presence of UM171*.** *HLF* expression is shown in single cell transcriptomes split up by treatment. Normalized expression data (z-score, after MAGIC imputation^49^) is expressed in color scale. ***F) Comparison of HLF expression specificity*** versus selected HSC-associated genes and common HSC surface marker genes. Gene-wise z-score distribution by treatment (fresh CD34+ and UM171 d7) is represented as density for each cell community (same color-code as in Fig. 1C). Mean z-score for HLF in HSPC cluster is provided as dotted reference line for each treatment. ***G-I) HLF expression is strongly enriched in HSC clusters in human bone marrow. G) Overview of cell clusters*.** (Human Cell Atlas; 101,935 cells; integrated data from eight donors, UMAP reduction; preprocessed data, clusters and labels adopted from^32^). ***H) HLF expression*.** (z-score normalized, after MAGIC imputation). ***I) Expression summary of selected HSC-associated genes in human bone marrow***. (scaled expression averaged for each hematopoietic cell population and donor (n=8) in the dataset, row-normalized color scale). ***J-L) HLF expression is restricted to HSC/MPP cluster in hematopoietic human fetal liver cells*.** ***J) Overview of cell communities*.** (Human Cell Atlas; 113,063 cells; integrated data from 14 fetal livers across four developmental stages, UMAP reduction; preprocessed data, clusters and labels adopted from^33^). Mac, macrophage; Neut-my, neutrophil–myeloid; Mono-mac, monocyte-macrophage; Early L/TL, early lymphoid/T lymphocyte; pro., progenitor. ***K) HLF* expression** (z-score normalized, after MAGIC imputation). ***L) Expression summary of HSC-associated genes* in fetal liver** (scaled expression was averaged for each hematopoietic cell population and four gestation stages (7-8, 9-11, 12-14 and 15-17 post-conception weeks), row-normalized color scale).

Next, we benchmarked the expression of *HLF* against (i) HSC-associated genes we and others identified (*AVP*^25^, *MLLT3*^26^) or that have previously been characterized in mice (*PRDM16*^27^, *GATA3*^8^, *HOXB5*^11^. *MEIS1*^12^, *MECOM*^14^, *FGD5*^9^ and alpha-Catulin (*CTNNAL1*)^10^), as well as against (ii) surface markers commonly used to prospectively isolate human HSCs (*CD34*^28^, CD201 (*PROCR*)^7^, CD49c (*ITGA3*)^21^, CD133 (*PROM1*)^29^, CD90 (*THY1*)^30^ and CD49f (*ITGA6*)^31^) (**Fig. 1F and Supplemental Fig. S2**). While varying degrees of HSPC-enriched expression were detectable for most of these genes (**pink density profiles in Fig. 1F and Supplemental Figure S2, Supplemental Tables T1 and T2**), *HLF, AVP, GATA3, MEIS1, HOXB5, MLLT3* and *MECOM* displayed the most specific expression within HSPCs of freshly purified human CD34^+^ cord blood cells (**Fig. 1F, left panels**). Of note, these genes generally performed better than HSC-associated surface antigens, consistent with the requirement to stain for several of these markers in combination to achieve high HSC enrichment. In seven-day UM171-supplemented cultures, *HLF, AVP, PRDM16* and *GATA3* exhibited the highest HSPC enrichment (**Fig. 1F, right panel**).

To extend our analysis to different developmental and physiological contexts, and to assess specificity in a larger diversity of hematopoietic lineages and intermediates, we examined public single cell transcriptomic datasets of adult bone marrow^32^ (**Fig. 1G-I and Supplemental Fig. S3A**) and fetal liver^33^ (**Fig. 1J-L and Supplemental Fig. S3B**), each aggregating more than 100,000 cells from several bio-informatically integrated specimens. These analyses revealed that *GATA3* is also expressed in innate-lymphoid cells (ILC), T-cells and NK cells and *PRDM16* expression in embryonic lymphoid/T-lymphoid precursors and NK cells, eliminating these genes from further consideration (**Fig. 1I and 1l and Supplemental Fig S3A and B**). Among hematopoietic cells in the adult bone marrow and fetal liver datasets, *HLF* exhibited pronounced HSPC-restricted expression and only negligible expression in a small subset of naïve T-cells in adult bone marrow (**Fig. 1H**). In non-hematopoietic cells, *HLF* expression was detectable in the stromal fraction of adult bone marrow, as well as in fetal liver fibroblasts and hepatocytes (**Fig. 1G-I**).

While *Arginine Vasopressin (AVP)* exhibited a similar expression profile as *HLF* in these analyses, we focused on *HLF* for downstream experiments based on its reported function in HSCs^34–36^.

In summary, *HLF* is a gene with highly selective expression in HSC-enriched sub-populations across their most relevant anatomical and ontogenetic sources. As such, it can be regarded as an attractive candidate gene to mark human hematopoietic stem cells independent of developmental or environmental (e.g. *ex vivo* culture) context.

### Engineering of a genomic *HLF*-reporter transgene in human cells

We reasoned that *HLF*-expression, if visualized genetically, could provide a specific readout to identify immature human blood cells. To this end, we devised a strategy to introduce a fluorescent reporter cassette into the endogenous *HLF* locus using nucleofection of a Cas9/sgRNA ribonucleoprotein complex and delivery of a homologous recombination (HR) template by recombinant adeno-associated virus (rAAV6) transduction^37^. To maintain HLF protein function, we targeted the 3’ end of the *HLF* open reading frame (**Fig. 2A and supplemental Fig. S4**). Several HR templates were designed to either knock-in an in-frame P2A-ZsGreen (ZsG) cassette or an IRES-ZsGreen cassette to capture endogenous *HLF* expression (**Supplemental Fig. S5**). Functional assessment in HLF-expressing HepG2 cells indicated that constructs with IRES-ZsGreen and either a P2A-linked Puromycin resistance or truncated EGFR^38^ yielded the highest reporter expression (**Supplemental Fig. S5D and E**).

**2).**
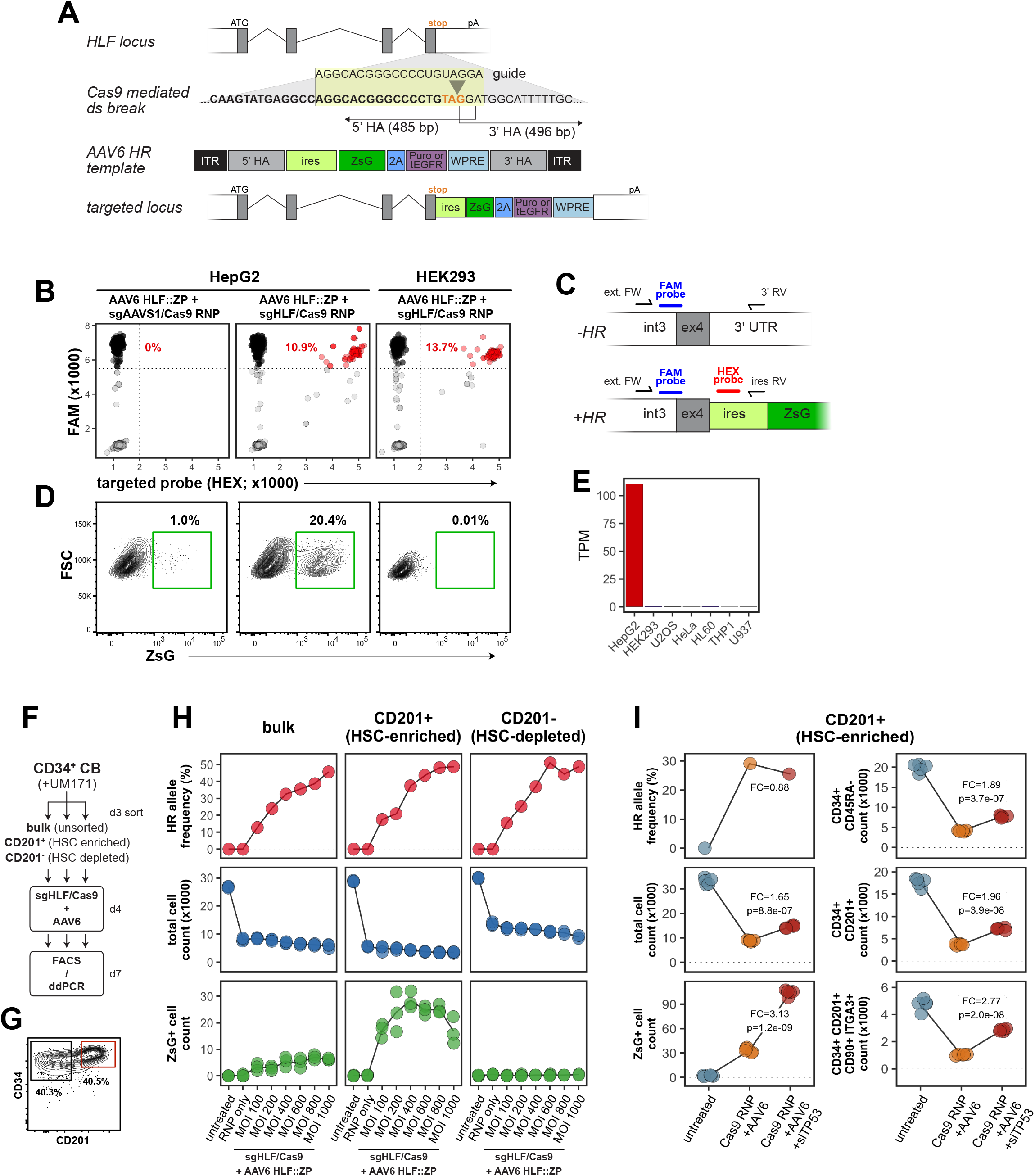
Engineering of a Human Genomic *HLF*-Reporter. ***A) Outline of the HLF-reporter targeting strategy using CRISPR/Cas9 and rAAV6*.** A sitespecific DSB at the HLF stop codon (orange) located in exon 4 is generated by a Cas9/sgHLF ribonucleoprotein (RNP) complex. This stimulates homologous recombination (HR) with a single-stranded donor template delivered through rAAV6 infection. The resulting HR event results in a transgenic locus that co-expresses the HLF open reading frame and a multifunctional ZsGreen (ZsG) expression cassette connected the endogenous *HLF* open reading frame by an EMCV internal ribosome entry site (ires). Grey boxes, HLF exons; white boxes, 5’ and 3’ untranslated regions; 3’/5’ HA, homology arms; purple box, puromycin resistance or truncated EGFR (tEGFR) sequence linked to ZsG by a P2A for optional drug or antibody mediated selection; WPRE, Woodchuck Hepatitis Virus Post-transcriptional Response Element; pA, endogenous *HLF* polyadenylation signal. ***B-E) Validation of the HLF reporter in human cell lines*.** HepG2 (HLF-expressing) and HEK293 (HLF non-expressing) cells were electroporated with Cas9/sgRNA RNP either as summarized in (A) or using sgAAVS1 as control. *HLF-ZP*, rAAV6 encoded HLF repair template driving expression of ZsG and Puromycin resistance. Representative data of two independent experiments. ***B*) Droplet digital PCR genotyping of targeted cell lines.** Black dots represent HR-negative and red dots represent HR-positive PCR droplets. HR percentages (printed in red) were calculated as HR-positive divided by the total number of specific amplicon-containing droplets (black and red). Representative data of two independent experiments. ***C*) ddPCR strategy.** ext. FW, external forward primer binding to a common region outside the 5’ HR; 3’ RV (reverse) primer amplifying unrecombined locus; ires RV (reverse) primer amplifying recombined locus; HR-negative and positive amplicons are detected by a common FAM-labeled probe and HR-positive amplicons are additionally recognized by a HEX-labeled probe that binds to the IRES region of the transgene. ***D) FACS analysis to detect reporter expression*.** ***E) HLF expression levels in selected cell lines*.** Data curated from Human Protein Atlas ^50^. ***F) Outline of experimental strategy to optimize reporter integration in CD34^+^ cord blood cells*.** ***G) FACS sorting strategy to enrich/deplete HSCs based on CD201 expression from expanded CD34^+^ cells at d3 of culture*.** ***H) Selective HLF-reporter expression in HSC-containing subfractions of CD34+ cord blood cell cultures*.** HR allele frequencies were determined from one of four replicate wells. Total and ZsG^+^ cell counts were determined by FACS on d7 and are normalized to 10e4 cells plated per 96-well post-electroporation at d4. MOI, multiplicity-of-infection. One of five independent experiments covering four biological replicates is shown. ***I) Effect of TP53 knock-down on HR and cell survival*.** CD201^+^ cells were sorted and targeted as in (h) (MOI400), and as an additional condition, electroporated with RNP and siRNA against TP53. One representative experiment of four independent experiments covering four biological replicates is shown.

Further validation of the reporter construct (**Fig. 2A**) in HepG2 and HEK293 cells revealed that only cognate pairing of guide RNA and repair template (**Fig. 2B**) resulted in targeting to the *HLF* locus, as demonstrated by droplet digital PCR (ddPCR) designed to detect the integrated but not the episomal reporter cassette (**Fig. 2C**). Importantly, targeted reporter integration resulted in stable ZsG expression in HepG2 but not in HEK293 cells (**Fig. 2D**), thus recapitulating endogenous *HLF* expression levels (**Fig. 2E**). These results not only provided proof-of-principle of the experimental approach but also demonstrated reporter functionality and selectivity.

We next assessed HR efficiency to the *HLF*-locus in cord blood derived HSPCs using reported settings^37^. To this end, we used a rAAV6 HR template which contained a constitutive Ubiquitin C (UbC) promoter driven Ametrine fluorescent protein cassette (**Supplemental Fig. S5A and S6A**). Since this promoter drives high expression after genomic integration but not from episomal rAAV vectors^39^, it provided a direct readout of insertion into the CRISPR-targeted *HLF*-locus. Indeed, we observed up to 55% of cells with high Ametrine expression in sgHLF/Cas9 RNP but not in mock electroporated cells indicative of targeted integration (**Supplemental Fig. S6B and S6C**). CD34 profiles were comparable between Ametrine positive and negative populations, suggesting largely uniform targeting efficiencies (**Supplemental Fig. S6B**). Notably, elevated rAAV6 concentrations resulted in marked cell toxicity, necessitating careful titration (**Supplemental Fig. S6C**).

We next tested the promoterless *HLF* reporter construct (**Fig. 2A and Supplemental Fig. S5D**) in CD34^+^ cells which, after 3 days of pre-expansion with UM171, were divided into HSC-enriched and depleted populations based on high or low CD201 surface expression, respectively^7^ (**Fig. 2F and G**). These sub-fractions, as well as unsorted (bulk) cells were expanded for an additional 24h, electroporated with Cas9/sgHLF RNP and transduced with the repair template encoding rAAV6 over a range of virus-to-cell ratios (multiplicity of infection, MOI100-1000). Finally, after three additional days of HSC-supportive culture, ddPCR and FACS analysis to evaluate HR efficiencies and reporter expression was performed (**Fig. 2F**).

Within corresponding experimental conditions, allelic HR frequencies were similar between bulk, HSC-enriched and depleted fractions and reached a maximum of ~50% with the highest tested rAAV6 virus titer of MOI1000 (**Fig. 2H top panels**). Strikingly, while HSC-depleted CD201-cultures gave rise to no ZsG^+^ cells (**Fig. 2H bottom right panel**) despite successful HR, reporter expressing cells were readily detectable in all other rAAV6-containing conditions (**Fig. 2H bottom panels**). In essence, pre-enrichment of CD201^+^ cells and rAAV6 transduction at MOI400 yielded the highest number of ZsG^+^ cells among all tested conditions (**Fig. 2H centre bottom panel**), although HR reached only intermediate levels with these parameters. This suggested rAAV6 toxicity for *HLF*-expressing cells at lower MOI compared to bulk cultures (**Fig. 2h middle panels and Supplemental Fig. S6C**).

Since Cas9/sgRNA electroporation impacted the overall cell recovery at day seven compared to untreated controls (**Fig. 2H middle panels**), we next tested whether transient p53 inhibition could partially alleviate this effect as suggested^40^.

Although co-electroporation of synthetic siRNA against p53 did not increase overall HR efficiency (**Fig. 2I, top left panel**), it resulted in more than threefold enhanced recovery of *HLF-ZsG* expressing cells (**Fig. 2I, left bottom panel**). While total cell numbers improved only marginally by transient p53 knock-down (**Fig. 2I, left middle panel**), a significantly more pronounced effect was detectable in immature HSPC subsets defined by surface marker expression (**Fig. 2I, right panels with increasing HSC enrichment from top to bottom**). These observations were in line with a report suggesting that nuclease-mediated gene editing results in p53-dependent proliferation arrest and functional impairment of the most immature HSPCs, and that transient p53 inhibition can partially overcome this effect^40^.

In summary, these experiments established a robust experimental approach to visualize endogenous *HLF*-expression through targeted genomic integration of a fluorescent reporter cassette in human cell lines and more importantly, in cord blood HSPCs.

### Selective *HLF*-transgene expression in immunophenotypic human LT-HSCs

Next, we aimed to validate expression of the *HLF-ZsG* reporter in the context of stem cell specific surface marker panels adapted for cultured cord blood cells^7,21^.

Using the established conditions described above, ~20% targeted allele frequencies were observed in CD201^-^ and CD201^+^ sorted cells (**Fig. 3A**), resulting in an average of 1.17% reporter expression specific to CD201^+^ sorted cells (**Fig. 3B and C**). Strikingly, reporter-positive cells largely expressed LT-HSC surface phenotypes defined by characteristic combinations of CD34, CD45RA, CD201, CD90 and ITGA3 (**Fig. 3d and e**). Dimensionality reduction of these surface marker and *HLF-ZsG* expression profiles indicated that *HLF-ZsG* expression defined a concise sub-population which clustered inside increasingly restricted immuno-phenotypically defined HSC populations (**Fig. 3F**). As expected, *HLF-ZsG* transgene expression increased within progressively restrictive HSC surface marker gates but only reached a maximum of 8% within the CD34^+^CD201^+^CD90^+^ITGA3^+^ population (**Fig. 3G and H**). Under consideration of the 20% targeted allele frequency, this suggested that HLF-expressing cells indeed represent only a fraction of this sub-population.

**3).**
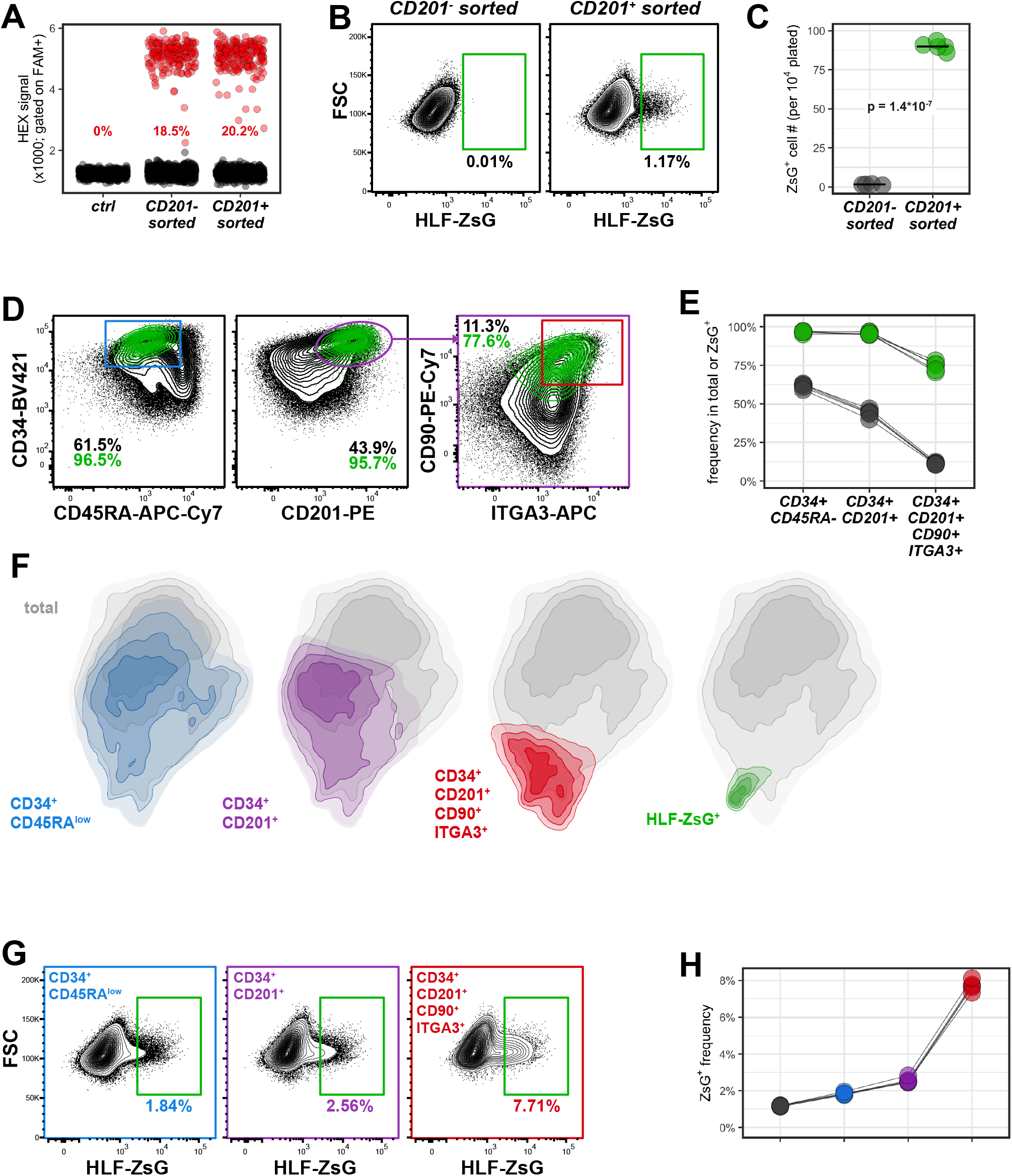
Selective HLF-Reporter Expression in Human Cord Blood Derived LT-HSC Populations. Cord blood derived CD34+ cells were processed as in **Fig. 2F** with the addition of siTP53 and transduction of rAAV6 *HLF-ZsG P2A tEGFR* at MOI400. One representative experiment of four independent experiments covering four biological replicates is shown. ***A) HR allele frequencies in CD201^+^/ pre-sorted fractions as determined by ddPCR*.** Gated on FAM+ (common probe) droplets, HEX+ droplets (red) identify HR allele amplicons. ***B-C) Reporter expression in CD201^+^/ pre-sorted fractions*.** Aggregated FACS analysis (b) and summary by repeat (n=4 for CD201- and n=5 for CD201+ sorted, unpaired two-sided t-test p-value is indicated) in (C). ***D-E) Immuno-phenotypes of ex vivo expanded (+UM171) HLF-targeted HSPCs*.** FACS analysis of total (black) versus reporter expressing (green) populations at day 7. Percentages of increasingly restricted HSC gates are provided for each population. Aggregated FACS data in (d) and summary by repeat in (e). ***F) Dimensional reduction based on FACS analysis*.** UMAP reduction using CD34, CD45RA, CD201, CD90, ITGA3 and ZsG FACS intensities from (d) was calculated and is represented as 2d density plot of all cells (grey, n = 306,797). Cells falling into HSC- or ZsG-gates are overlayed and color-coded as in (d). ***G-H) HLF-reporter expression in immuno-phenotypic HSC gates*.** Reverse gating of the same data as above showing reporter expression in increasingly restricted HSC gates. Aggregated FACS data from all repeats in (G), summary in (H).

These results suggested that *HLF-ZsG* expression *per se* has the potential to surrogate multiparametric FACS analysis to identify the most immature cells in *ex vivo* cord blood cell cultures.

### *HLF* expression identifies repopulating cells in CD34^+^ cord blood cultures

To test the ability of the reporter to identify functional HSCs, we sorted and transplanted either bulk *HLF-ZsG* targeted (**Fig. 4A, black colour code**), reporter expressing (**Fig. 4A, green**) or non-expressing sub-populations (**Fig. 4A, blue**) of CD201 pre-enriched HSPCs. To control for adverse effects of the targeting procedure, two additional cohorts received either untreated parental cells (**Fig. 4A, red**) or cells electroporated with a neutral guide RNA (sgAAVS1) (**Fig. 4A, purple**). Importantly, transplanted cell doses were proportional of reporter positive versus negative subsets in the total targeted population (**summarized in Fig. 4B**).

**4).**
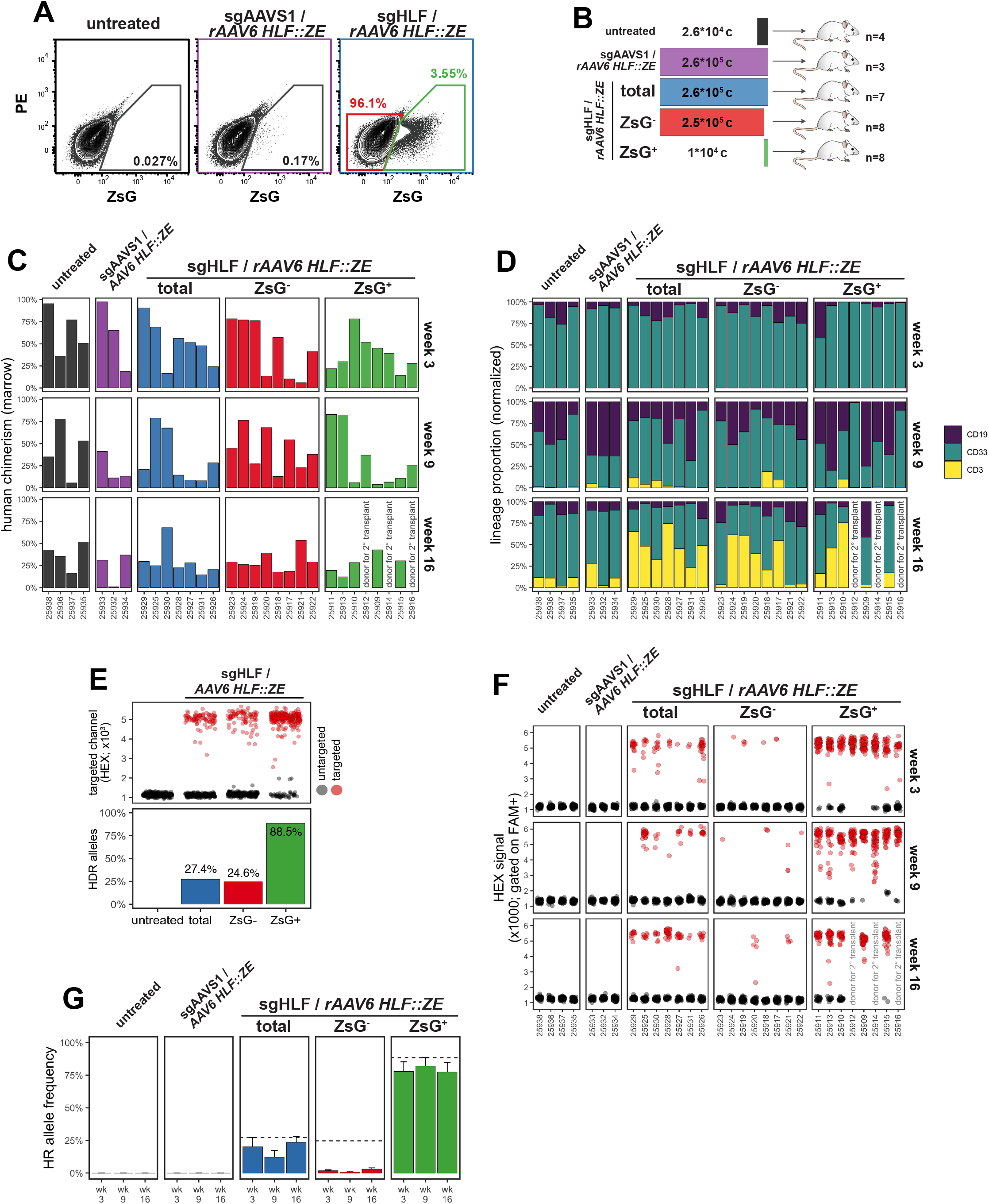
HLF-Reporter Labels Repopulating Cells in CD34+ Cord Blood Cell Cultures. ***A) FACS plots showing the sorting of HLF-reporter targeted population for transplantation*.** *rAAV6 HLF-ZE*: recombinant rAAV6 particle encoding an *HLF* repair template with ires ZsGreen P2A tEGFR cassette. A pool of eight cord blood units was split into three and processed as indicated. ***B) Summary of transplantation layout*.** Transplantation cohorts and cell doses are represented using the same color-code as in (***A***). ***C) Human engraftment summary of transplanted NSGS recipients*.** Human bone marrow chimerism determined based on human CD45+ cells among total CD45+ (mouse and human) cells at short (week 3), intermediate (week 9) and long-term (week 16) post-transplantation timepoints is plotted using the same color code as in (a) and (b). Each recipient mouse is represented along the x-axis (NSGS-ID). Recipients are arranged by descending average reconstitution across all timepoints. Recipients #25912, #25914 and #25916 were sacrificed at week 10 post-transplantation to be used as donors for secondary transplantation (summarized in **Fig. 5**). ***D) Lineage proportion of transplanted recipients*.** Bone marrow biopsies were analyzed and are arranged along timepoints and the individual recipients as in (e). Normalized proportions of B-cells (CD19), myeloid cells (CD33) and T-cells (CD3) within human CD45^+^ cells for each timepoint and recipient are color-coded as indicated. ***E) HR allele frequencies in pre-transplanted cell populations*.** top panel: ddPCR droplets are pre-gated based on FAM-positivity, black droplets represent FAM+/HEX-events indicative of untargeted alleles, red droplets (FAM/HEX double positive) indicate targeted alleles, sub-sampled to 300 droplets per specimen. bottom panel, quantification summary of HR frequencies calculated based on targeted/(untargeted+targeted) droplets. ***F) ddPCR analysis of bone marrow biopsies at weeks 3, 9 and 16*.** Specimens are arranged as in (C), ddPCR droplets are represented as in (E), sub-sampled to 50 droplets per specimen and timepoint. ***G) HR allele tracing summary*.** Summarized data representation of (E) and (F). Dashed red lines represent allele frequencies at time of transplantation. Bars represent average HR allele frequencies from (e) with standard error bars, color-codes as in (A-C). One representative experiment of two independent experiments is summarized.

We assessed the engraftment potential of these cells in transplanted NSGS recipients at short (3 weeks), intermediate (9 weeks) and long-term (16 weeks) timepoints. Even though at least 25-fold fewer *HLF-ZsG* expressing cells were transplanted compared to reporter non-expressing or total *HLF-ZsG* targeted cells, comparable reconstitution levels (**Fig. 4C**) and similar lineage contributions (**Fig. 4D**) were achieved in these cohorts. Furthermore, we observed similar levels of human hematopoietic chimerism between sgAAVS1 controls and *HLF-ZsG* targeted recipients, indicating that genetic manipulation of the *HLF* locus has little functional impact on HSC activity beyond what can be attributed to RNP electroporation and rAAV6 transduction (**Fig. 4C first three panel columns**).

While these results suggested a strong enrichment for reconstitution activity in reporter expressing cells based on transplanted cell doses, we additionally traced the engineered reporter allele in transplanted recipients by ddPCR to test whether human engraftment in the reporter non-expressing cohort had mainly emerged from non-targeted HSCs. Indeed, this was confirmed by the observation that HR allele ratios of 24.6% in the reporter non-expressing fraction at transplantation (**Fig. 4E, blue**) dropped sharply in the progeny of these cells *in vivo* (**Fig. 4F and G**). On the contrary, HR allele frequencies remained largely stable between pre- and post-transplantation timepoints in both total targeted and sorted reporter positive populations. Strikingly, HR allele frequency reached 88.5% in the *HLF-ZsG* expressing fraction indicating that most of these cells carried bi-allelic reporter integration (**Fig. 4E**).

Taken together, these results demonstrated that reporter-visualized *HLF*-expression is able to identify multipotent cells with high *in vivo* regenerative potential in CD34^+^ cord blood cell cultures.

### *HLF*-expression labels HSCs with extensive self-renewal capacity

At three weeks post-transplantation, a well-defined reporter positive sub-fraction within CD34-high bone marrow cells was detectable in recipients transplanted with total or ZsG-positive *HLF-ZsG* targeted cells but was entirely absent in ZsG-negative recipients (**Fig. 5A**). To test whether *HLF* expression continued to label stem cells after transplantation, we selected three *HLF-ZsG* targeted ZsG^+^ primary recipients with detectable ZsG^+^ cells at week 10.5 (**Fig. 5B**) and high (>96% each) HR allele ratios (**Fig. 4F**) as donors for secondary transplantation. We first magnetically enriched CD34^+^ cells from the pooled bone marrow of these donors and then sorted reporter-expressing cells as well as a corresponding population that expressed similar CD34 levels but was 16.6-fold more abundant. Corresponding cell numbers of these populations (900 and 1.5*10e4, respectively) were intra-hepatically transplanted into newborn secondary recipient mice (**Fig. 5C**). Significant differences in human engraftment between HLF-ZsG^+^ and CD34^high^/HLF-ZsG^-^ secondary recipients were observed at short-term, intermediate and long-term post-transplantation timepoints (**Fig. 5D**). More specifically, while all HLF-ZsG^-^ secondary recipients were characterized by low-level and transient reconstitution, pronounced and multi-lineage human chimerism was detectable in at least four of ten HLF-ZsG^+^ recipients as long as 16 weeks post-transplantation (**Fig. 5D and E**).

**5).**
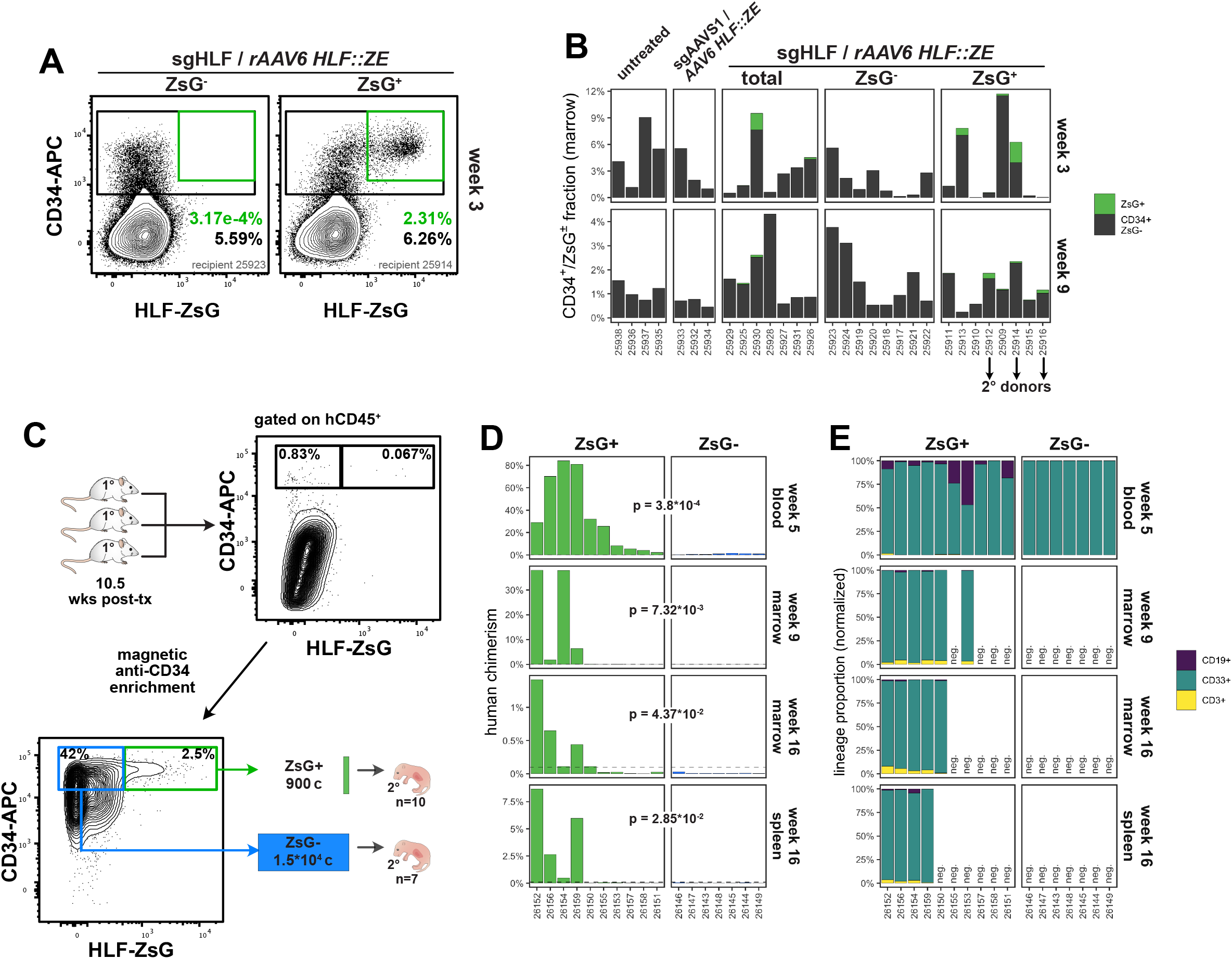
HLF-Reporter Labels Human HSCs with Extensive Self-Renewal Capacity. ***A) FACS plots of CD34+/HLF-ZsG+ population***. Representative bone marrow biopsies of reporter-negative (*sgHLF/ rAAV6 HLF-ZE* targeted, ZsG^-^ sorted, left) and reporter-positive (*sgHLF/ rAAV6 HLF-ZE* targeted, ZsG^+^ sorted, right panel) primary recipients, gated on human CD45+. ***B) Summary of CD34+/HLF-ZsG+ population*.** Population overview of all primary recipients, pre-gated on human CD45+, recipient mice are arranged according to engraftment levels as in Fig. 4c. ***C) Strategy for secondary transplantation*.** Bone marrow of three primary recipients (*sgHLF/ rAAV6 HLF-ZE* targeted, ZsG^+^ sorted cohort) was pooled and magnetically enriched for human CD34 expression. Reporter-expressing (ZsG+) and non-expressing cells (ZsG-) with comparable levels of CD34 expression were sorted for transplantation. Intra-hepatic transplantation into newborn NSGS recipients as outlined. Corresponding cell doses of HLF-ZsG+ (n=10) and HLF-ZsG- (n=7) were transplanted. ***D) Human engraftment summary of secondary recipients*.** Human bone chimerism in indicated tissues was determined based on human CD45-expressing cells among total (mouse and human) CD45+ cells at short (week 5, blood), intermediate (week 9, marrow) and long-term (week 16, marrow and spleen) post-transplantation. Dashed line represents the 0.1% mark used as cut-off for engraftment positivity. Significance was calculated by unpaired, one-sided (alternative = “greater”) Wilcoxon test and is provided as p-value for a given comparison. ***E) Lineage output of engrafted human cells*.** Positive specimens from (D) are shown and color-coded for B-cells (CD19+), myeloid cells (CD33+) and T-cells (CD3). Normalized for lineage proportions within human CD45+ cells. Samples with less than 0.1% of human chimerism are designated negative (neg.).

Based on these observations, we conclude that in addition to labeling human HSCs *ex vivo, HLF* expression continues to mark hematopoietic stem cells with extensive reconstitution activity *in vivo*. These results thus demonstrate the potential of *HLF* reporter transgenesis to visualize human blood stem cells in real-time under experimental conditions.

## Discussion

With the aim to develop an approach to directly mark the rare stem cell fraction within the human blood system, we identify *HLF* as one of the most selectively expressed gene in human HSCs, corroborating its reported roles in the transcriptional regulation of HSC self-renewal and multipotency in mice^34,35^. Interestingly, in addition to these endogenous activities, ectopically expressed HLF can impart self-renewal to differentiation-committed blood cells through its DNA binding activity, either in the context of recurrent chromosomal fusions with *TCF3* in t(17;19) acute B-lymphoblastic leukemia^41^ or as a reprogramming factor in murine induced hematopoietic stem cells^42^, suggesting a role as a central master transcription factor in HSCs. Transgenic labeling of mouse HSCs has only recently been used to study these cells in their physiological and anatomical contexts during development^13^ and in the adult^10–12,14^.

With this study, we provide the first demonstration of transgenic labeling of human HSCs to date. Our approach builds directly on recently established methodology allowing precise genetic manipulations in human HSCs *ex vivo*^43^. Nonetheless, a number of challenges that equally affect therapeutic and experimental gene editing in HSCs remain. These challenges are primarily connected to the functional impact of both the delivery (e.g. electroporation, immune response to HR templates and sgRNA)^44,45^ and activities of the editing machinery (DNA double strand breaks)^40,46^ and represent important areas of investigation to optimize targeted gene therapies in stem cells.

An additional limitation of the presented strategy is that a significant fraction of HSCs remains untargeted and therefore escapes labeling. Introduction of a constitutive marker cassette in the HR template has the potential to circumvent this shortcoming but poses the risk of transcriptional interference with the targeted gene, as observed (**Supplemental Fig. S5**).

Notwithstanding these limitations, we provide a directly quantifiable platform to optimize gene editing in human HSCs under function-preserving conditions. Moreover, quantitative readout of the human HSC reporter will likely find utility in efforts to further optimize human HSC expansion conditions, either through screening for pharmacological self-renewal agonists or through systematic optimization of media composition and overall culture design.

## Acknowledgments

We thank Keith Humphries, Julie Lessard and Trang Hoang for critical reading of the manuscript, Annie Gosselin and Angelique Bellemare for assistance in FACS sorting, Melanie Frechette and Valérie Blouin-Chagnon for assistance with mouse experiments, and Mike Tyers for providing access to nucleofection equipment.

## Author contributions

B.L.: project conception, designed and performed all experiments, experimental and bioinformatic data analyses and interpretation, generated all figures, wrote manuscript; J.C. and E.T.: technical setup and assistance with CD34+ cell culture and FACS experiments, interpretation of results; T.M. designed and assisted with ddPCR analyses; S.C., I.B.: assistance with CD34+ cell purification and banking; N.M.: assistance with CD34+ cell purification, banking and mouse experiments; G.S.: project supervision and coordination, experimental design, interpretation of results and manuscript preparation.

## Citations

1. Till JE, McCulloch EA. A Direct Measurement of the Radiation Sensitivity of Normal Mouse Bone Marrow Cells. Radiat. Res. 1961;14(2):213.

2. Siminovitch L, McCulloch EA, Till JE. The distribution of colony-forming cells among spleen colonies. J Cell Comp Physiol. 1963;62(3):327–336.

3. Wu AM, Till JE, Siminovitch L, McCulloch EA. A cytological study of the capacity for differentiation of normal hemopoietic colony-forming cells. Journal of Cellular Physiology. 1967;69(2):177–184.

4. Wilkinson AC, Igarashi KJ, Nakauchi H. Haematopoietic stem cell self-renewal in vivo and ex vivo. Nat Rev Genet. 2020;132:1–14.

5. Doulatov S, Notta F, Laurenti E, Dick JE. Hematopoiesis: A Human Perspective. Cell Stem Cell. 2012;10(2):120–136.

6. Dorrell C, Gan OI, Pereira DS, Hawley RG, Dick JE. Expansion of human cord blood CD34+CD38-cells in ex vivo culture during retroviral transduction without a corresponding increase in SCID repopulating cell (SRC) frequency: dissociation of SRC phenotype and function. Blood. 2000;95(1):102–110.

7. Fares I, Chagraoui J, Lehnertz B, et al. EPCR expression marks UM171-expanded CD34+ cord blood stem cells. Blood. 2017;129(25):3344–3351.

8. Frelin C, Herrington R, Janmohamed S, et al. GATA-3 regulates the self-renewal of long-term hematopoietic stem cells. Nat Immunol. 2013;14(10):1037–1044.

9. Gazit R, Mandal PK, Ebina W, et al. Fgd5 identifies hematopoietic stem cells in the murine bone marrow. J Exp Med. 2014;211(7):1315–1331.

10. Acar M, Kocherlakota KS, Murphy MM, et al. Deep imaging of bone marrow shows non-dividing stem cells are mainly perisinusoidal. Nature. 2015;526(7571):126–130.

11. Chen JY, Miyanishi M, Wang SK, et al. Hoxb5 marks long-term haematopoietic stem cells and reveals a homogenous perivascular niche. Nature. 2016;530(7589):223–227.

12. Xiang P, Wei W, Hofs N, et al. A knock-in mouse strain facilitates dynamic tracking and enrichment of MEIS1. Blood Advances. 2017;1(24):2225–2235.

13. Yokomizo T, Watanabe N, Umemoto T, et al. Hlf marks the developmental pathway for hematopoietic stem cells but not for erythro-myeloid progenitors. J Exp Med. 2019;280(2):jem.20181399-320.

14. Christodoulou C, Spencer JA, Yeh S-CA, et al. Live-animal imaging of native haematopoietic stem and progenitor cells. Nature. 2020;514:1–6.

15. CRISPR/Cas9 β-globin gene targeting in human haematopoietic stem cells. Nature. 2016;539(7629):384–389.

16. Boitano AE, Wang J, Romeo R, et al. Aryl hydrocarbon receptor antagonists promote the expansion of human hematopoietic stem cells. Science. 2010;329(5997):1345–1348.

17. Fares I, Chagraoui J, Gareau Y, et al. Pyrimidoindole derivatives are agonists of human hematopoietic stem cell self-renewal. Science. 2014;345(6203):1509–1512.

18. Chaurasia P, Gajzer DC, Schaniel C, D’Souza S, Hoffman R. Epigenetic reprogramming induces the expansion of cord blood stem cells. J Clin Invest. 2014;124(6):2378–2395.

19. Mantel CR, O’Leary HA, Chitteti BR, et al. Enhancing Hematopoietic Stem Cell Transplantation Efficacy by Mitigating Oxygen Shock. Cell. 2015;161(7):1553–1565.

20. Bai T, Li J, Sinclair A, et al. Expansion of primitive human hematopoietic stem cells by culture in a zwitterionic hydrogel. Nat Med. 2019;25(10):1566–1575.

21. Tomellini E, Fares I, Lehnertz B, et al. Integrin-α3 Is a Functional Marker of Ex Vivo Expanded Human Long-Term Hematopoietic Stem Cells. Cell Reports. 2019;28(4):1063–1073.e5.

22. Uhlen M, Karlsson MJ, Zhong W, et al. A genome-wide transcriptomic analysis of protein-coding genes in human blood cells. Science. 2019;366(6472):eaax9198.

23. Velten L, Haas SF, Raffel S, et al. Human haematopoietic stem cell lineage commitment is a continuous process. Nat Cell Biol. 2017;19(4):271–281.

24. Haas S, Trumpp A, Milsom MD. Causes and Consequences of Hematopoietic Stem Cell Heterogeneity. Cell Stem Cell. 2018;22(5):627–638.

25. Zheng S, Papalexi E, Butler A, Stephenson W, Satija R. Molecular transitions in early progenitors during human cord blood hematopoiesis. Molecular Systems Biology. 2018;14(3):e8041.

26. Calvanese V, Nguyen AT, Bolan TJ, et al. MLLT3 governs human haematopoietic stem-cell self-renewal and engraftment. Nature. 2019;576(7786):1–6.

27. Aguilo F, Avagyan S, Labar A, et al. Prdm16 is a physiologic regulator of hematopoietic stem cells. Blood. 2011;117(19):5057–5066.

28. Civin CI, Strauss LC, Brovall C, et al. Antigenic analysis of hematopoiesis. III. A hematopoietic progenitor cell surface antigen defined by a monoclonal antibody raised against KG-1a cells. J Immunol. 1984;133(1):157–165.

29. Yin AH, Miraglia S, Zanjani ED, et al. AC133, a Novel Marker for Human Hematopoietic Stem and Progenitor Cells. Blood. 1997;90(12):5002–5012.

30. Baum CM, Weissman IL, Tsukamoto AS, Buckle AM, Peault B. Isolation of a candidate human hematopoietic stem-cell population. Proc Natl Acad Sci USA. 1992;89(7):2804–2808.

31. Notta F, Doulatov S, Laurenti E, et al. Isolation of Single Human Hematopoietic Stem Cells Capable of Long-Term Multilineage Engraftment. Science. 2011;333(6039):218–221.

32. Hay SB, Ferchen K, Chetal K, Grimes HL, Salomonis N. The Human Cell Atlas bone marrow single-cell interactive web portal. Exp Hematol. 2018;68:51–61.

33. Popescu D-M, Botting RA, Stephenson E, et al. Decoding human fetal liver haematopoiesis. Nature. 2019;574(7778):365–371.

34. Komorowska K, Doyle A, Wahlestedt M, et al. Hepatic Leukemia Factor Maintains Quiescence of Hematopoietic Stem Cells and Protects the Stem Cell Pool during Regeneration. Cell Reports. 2017;21(12):3514–3523.

35. Wahlestedt M, Ladopoulos V, Hidalgo I, et al. Critical Modulation of Hematopoietic Lineage Fate by Hepatic Leukemia Factor. Cell Reports. 2017;21(8):2251–2263.

36. Garg S, Reyes-Palomares A, He L, et al. Hepatic leukemia factor is a novel leukemic stem cell regulator in DNMT3A, NPM1, and FLT3-ITD triple-mutated AML. Blood. 2019;134(3):263–276.

37. Bak RO, Dever DP, Porteus MH. CRISPR/Cas9 genome editing in human hematopoietic stem cells. Nature protocols. 2018;13(2):358–376.

38. Wang X, Chang WC, Wong CW, et al. A transgene-encoded cell surface polypeptide for selection, in vivo tracking, and ablation of engineered cells. Blood. 2011;118(5):1255–1263.

39. Charlesworth CT, Camarena J, Cromer MK, et al. Priming Human Repopulating Hematopoietic Stem and Progenitor Cells for Cas9/sgRNA Gene Targeting. Mol Ther Nucleic Acids. 2018;12:89–104.

40. Schiroli G, Conti A, Ferrari S, et al. Precise Gene Editing Preserves Hematopoietic Stem Cell Function following Transient p53-Mediated DNA Damage Response. Cell Stem Cell. 2019;24(4):551–565.e8.

41. Inaba T, Roberts WM, Shapiro LH, et al. Fusion of the leucine zipper gene HLF to the E2A gene in human acute B-lineage leukemia. Science. 1992;257(5069):531–534.

42. Riddell J, Gazit R, Garrison BS, et al. Reprogramming committed murine blood cells to induced hematopoietic stem cells with defined factors. Cell. 2014;157(3):549–564.

43. Dever DP, Bak RO, Reinisch A, et al. CRISPR/Cas9 β-globin gene targeting in human haematopoietic stem cells. Nature. 2016;539(7629):384–389.

44. Cromer MK, Vaidyanathan S, Ryan DE, et al. Global Transcriptional Response to CRISPR/Cas9-AAV6-Based Genome Editing in CD34+ Hematopoietic Stem and Progenitor Cells. Mol Ther. 2018;26(10):2431–2442.

45. van Haasteren J, Li J, Scheideler OJ, Murthy N, Schaffer DV. The delivery challenge: fulfilling the promise of therapeutic genome editing. Nature Biotechnology. 2020;38(7):845–855.

46. Ferrari S, Jacob A, Beretta S, et al. Efficient gene editing of human long-term hematopoietic stem cells validated by clonal tracking. Nature Biotechnology. 2020;83:1–11.

47. Stuart T, Butler A, Hoffman P, et al. Comprehensive Integration of Single-Cell Data. Cell. 2019;177(7):1888–1902.e21.

48. Weinreb C, Wolock S, Klein AM. SPRING: a kinetic interface for visualizing high dimensional single-cell expression data. Bioinformatics. 2018;34(7):1246–1248.

49. van Dijk D, Sharma R, Nainys J, et al. Recovering Gene Interactions from Single-Cell Data Using Data Diffusion. Cell. 2018;174(3):716–729.e27.

50. Thul PJ, Åkesson L, Wiking M, et al. A subcellular map of the human proteome. Science. 2017;356(6340):eaal3321.

