## Supplementary figures and images for "*HLF* Expression Defines the Human Hematopoietic Stem Cell State"

### supplemental Figure 1

**Supplemental Fig. S1:** Intersection of EPCR and ITGA3 transcriptome datasets.

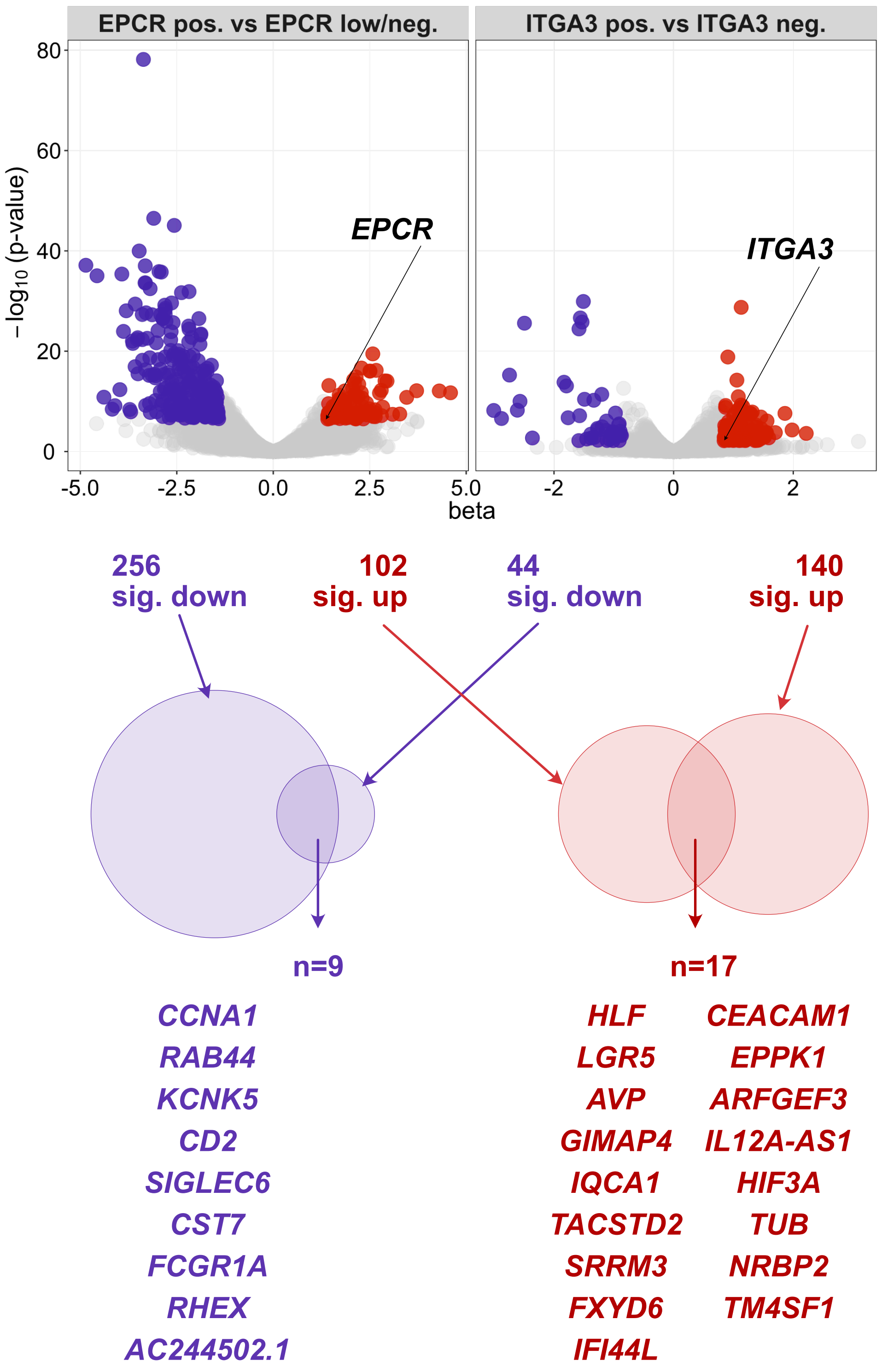

### supplemental Figure 5

#### <u>Supplemental Fig. S5:</u> Validation of *HLF*-reporter targeting constructs in HepG2 cells.

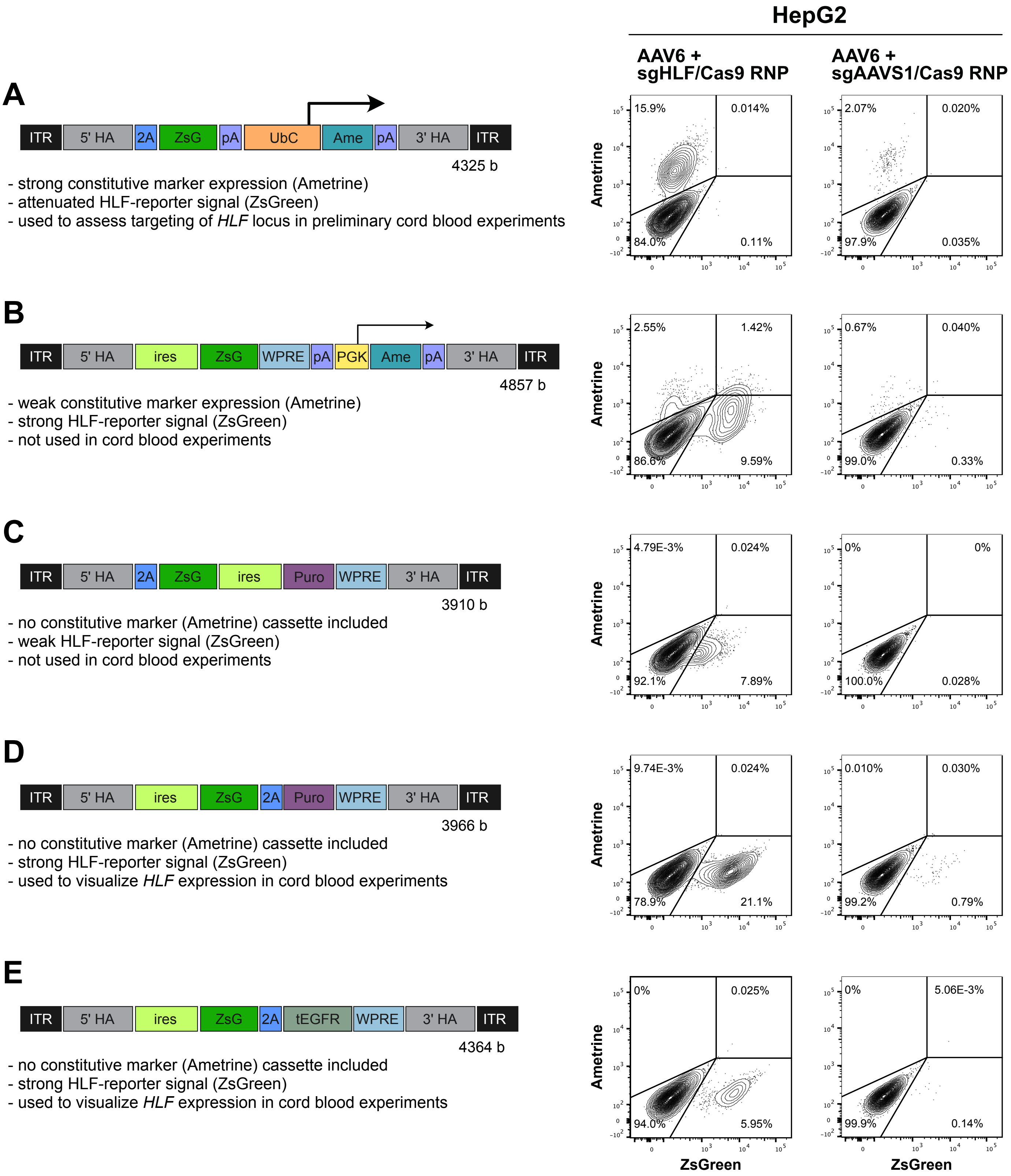

### supplemental Figure 6

#### Supplemental Fig. S6: Validation of rAAV6 HR at the *HLF*-target site in CD34+ cord blood cells.

A

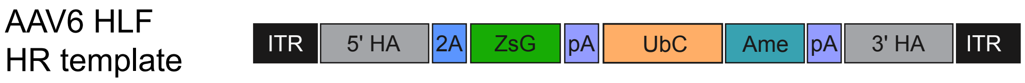


B

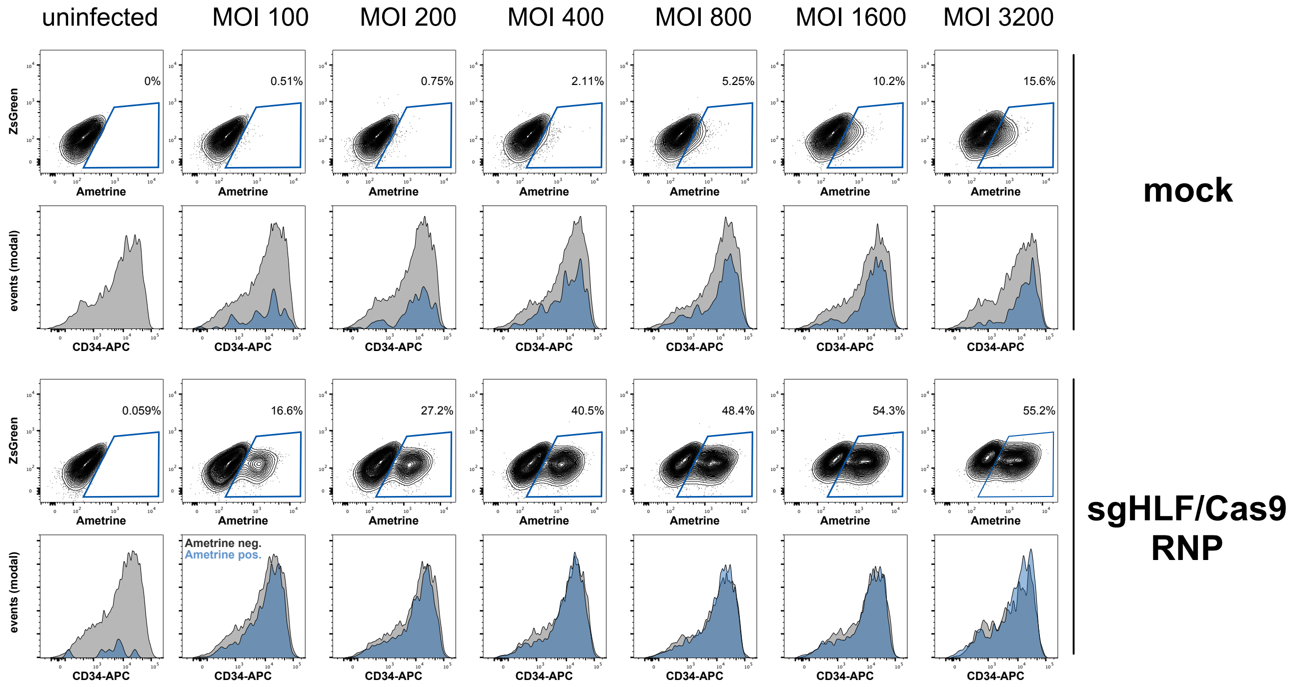


C

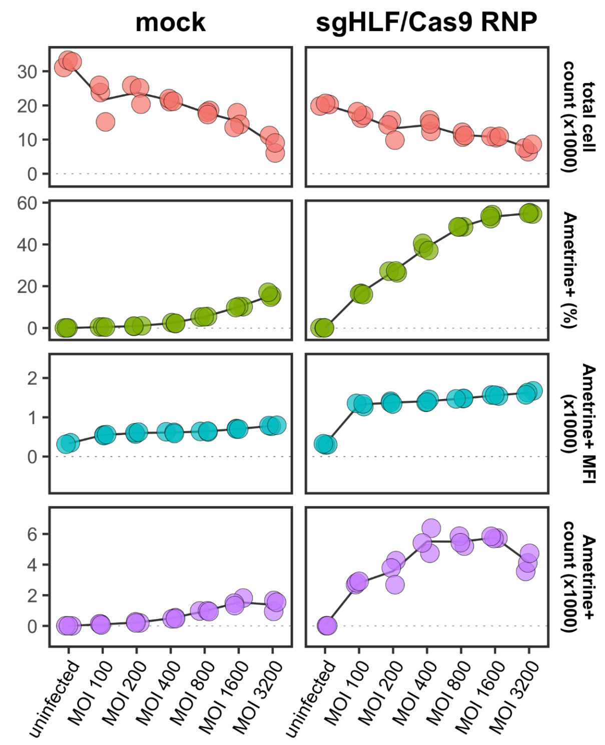
