## supplemental Figure 2 for "*HLF* Expression Defines the Human Hematopoietic Stem Cell State"

**<u>Supplemental Fig. S2:</u> Expression of HSC-associated genes and surface markers in fresh and UM171-expanded CD34+ cord blood cells.**

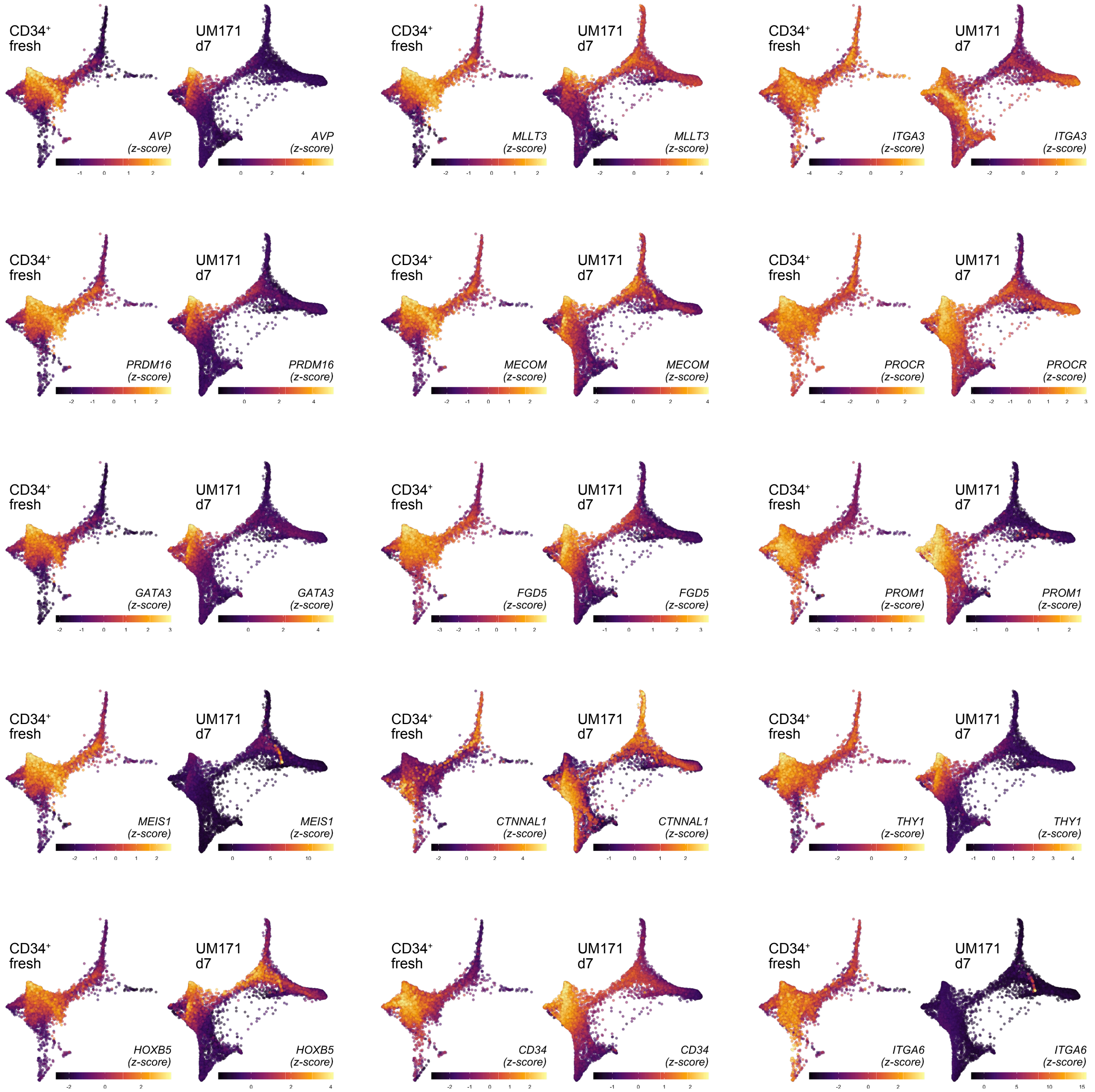
