## supplemental Figure 3 for "*HLF* Expression Defines the Human Hematopoietic Stem Cell State"

**<u>Supplemental Fig. S3:</u> Expression of HSC-associated genes and surface markers in human bone marrow and fetal liver.**

### a) Human Cell Atlas bone marrow

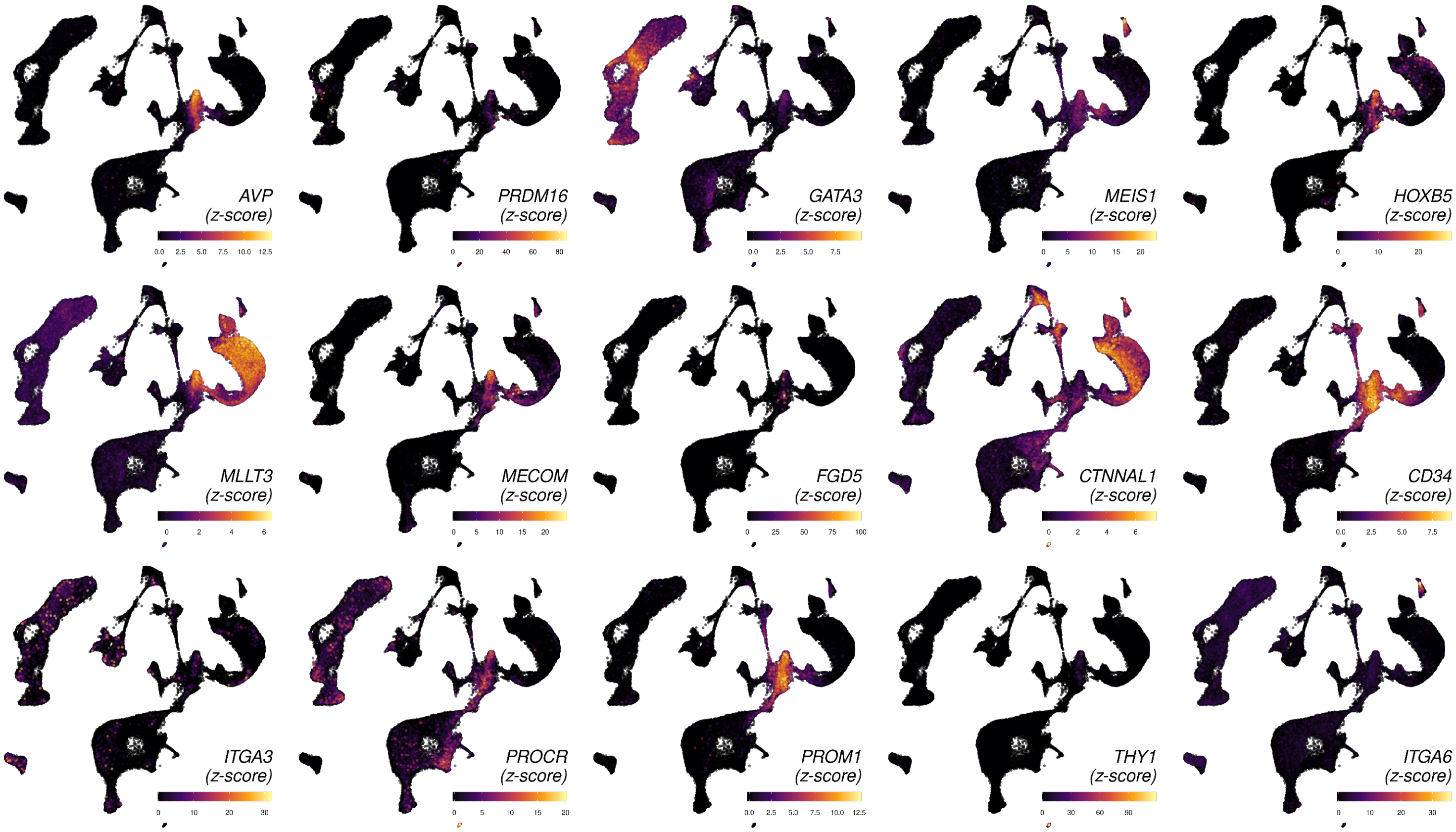


### b) Human Cell Atlas fetal liver

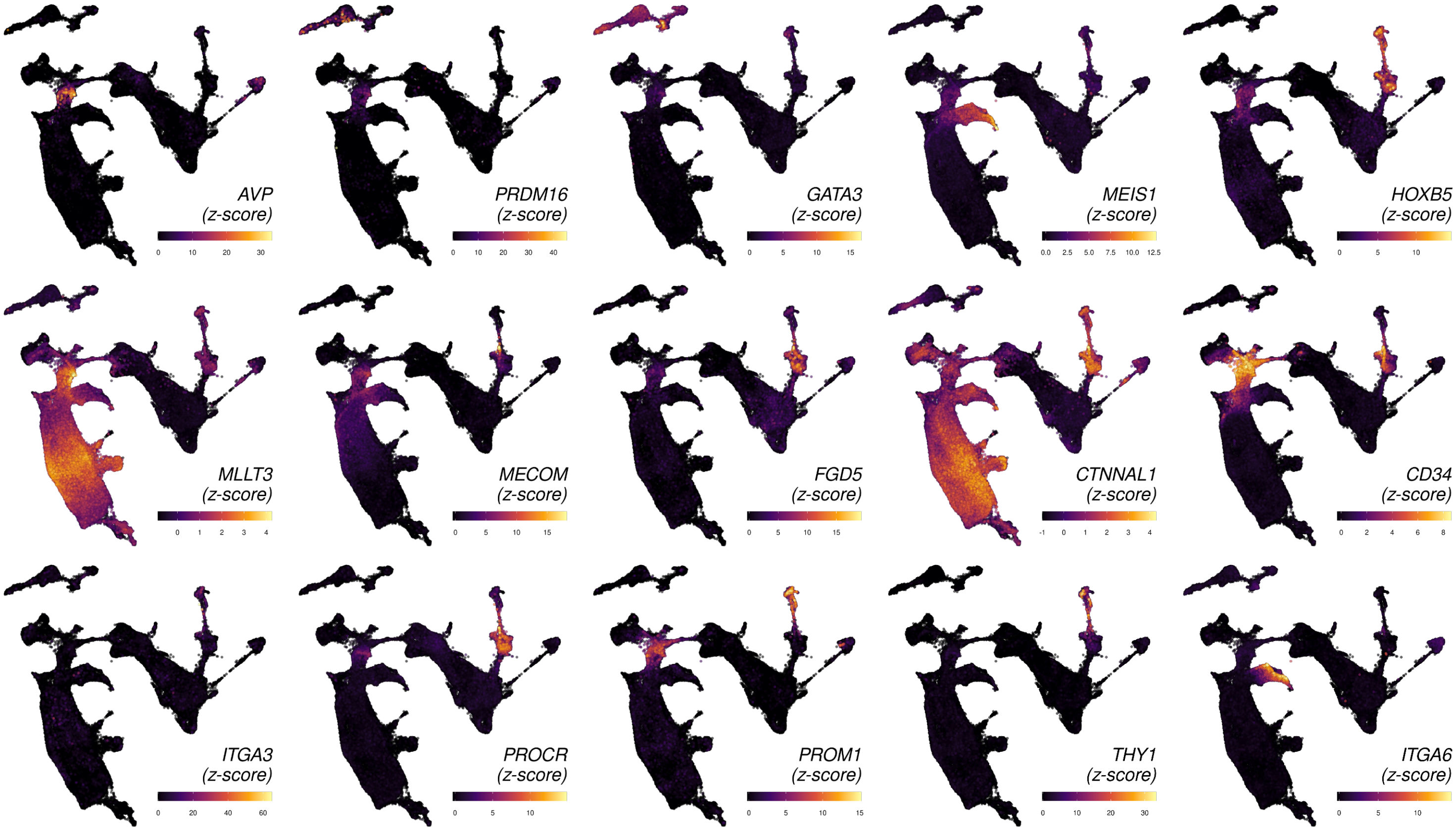
