## supplemental Figure 4 for "*HLF* Expression Defines the Human Hematopoietic Stem Cell State"

**<u>Extended Data Fig. S4:</u> High efficiency of Cas9/sgHLF RNP mediated Indel formation at the target site.**

**a**

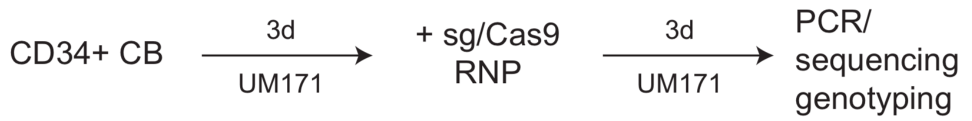


**b**

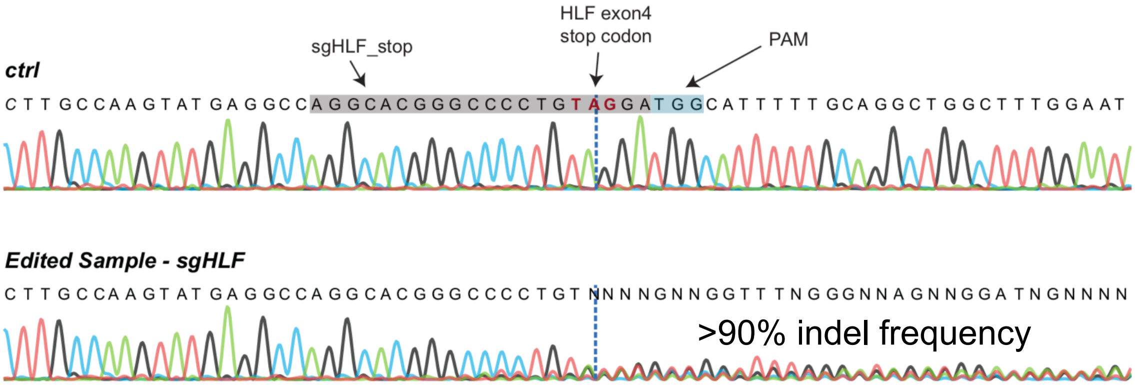
