## supplemental Figure 7 for "*HLF* Expression Defines the Human Hematopoietic Stem Cell State"

**Supplemental Fig. E7:** **Gating strategies to assess human reconstitution in transplanted hosts.**

**A) primary recipients; bone marrow aspiration, early timepoint (3 weeks)**

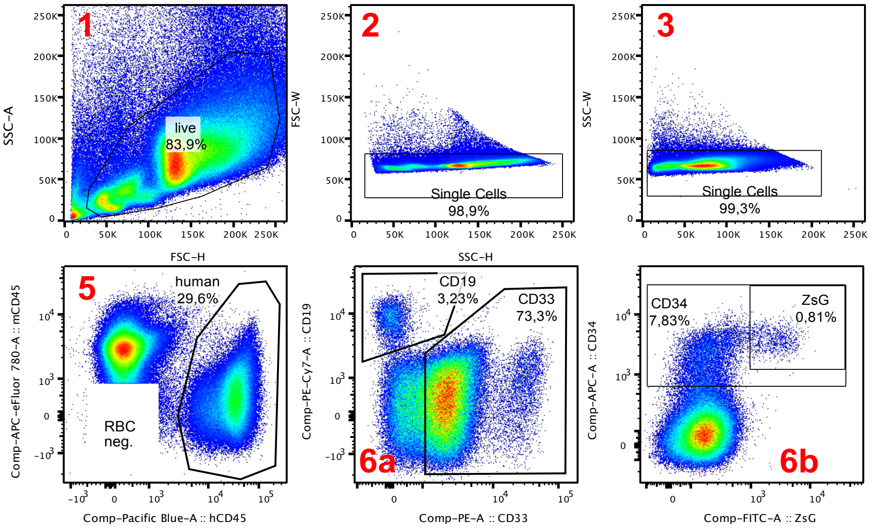


**B) primary recipients; bone marrow aspiration, endpoint (16 weeks)**

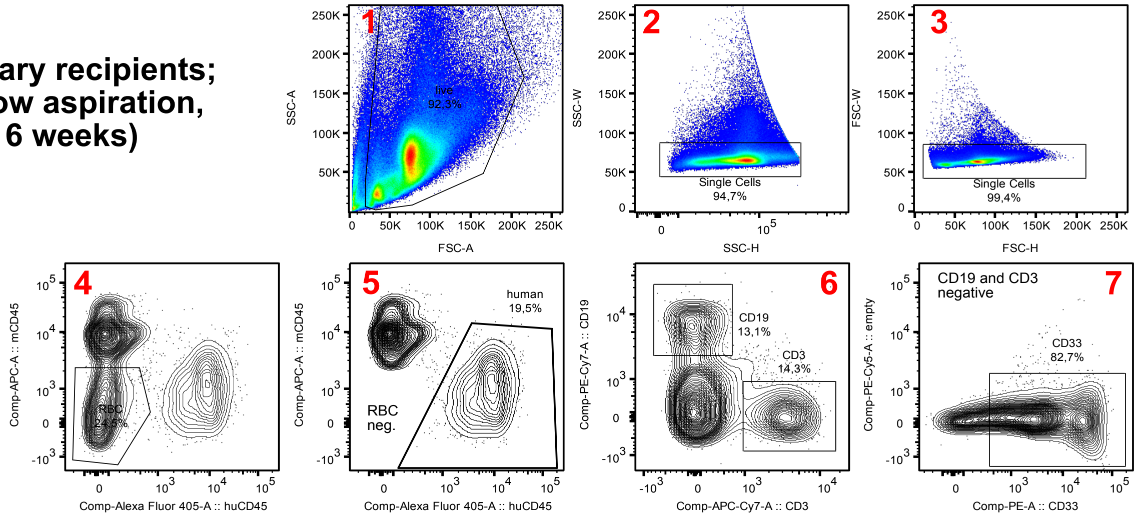


**C) secondary recipients; bone marrow aspiration, intermediate timepoint (9 weeks)**

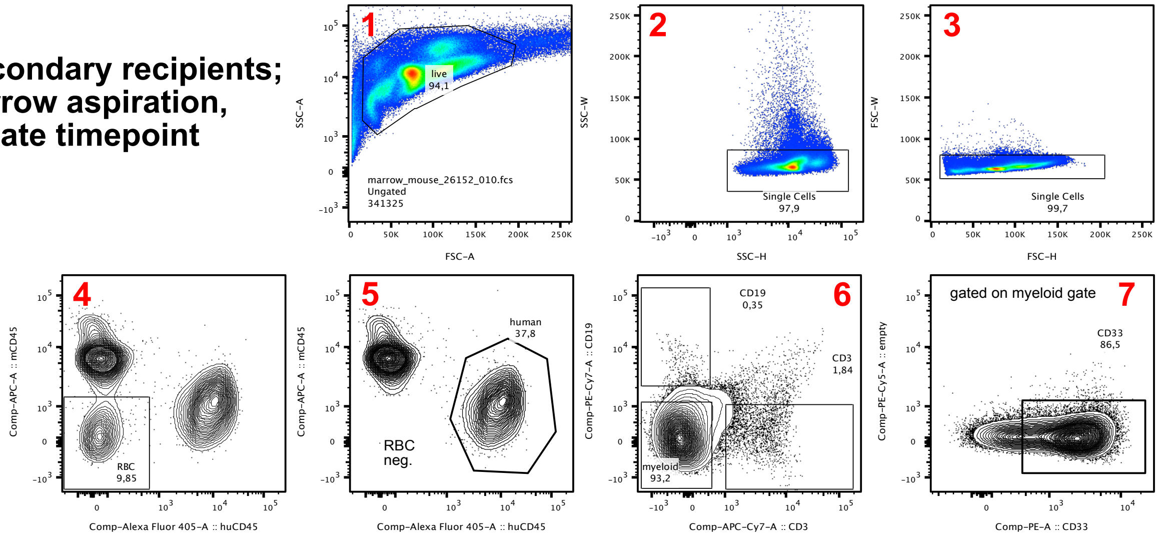
