## supplemental Figure 8 for "*HLF* Expression Defines the Human Hematopoietic Stem Cell State"

**<u>Supplemental Fig. E8:</u> Gating strategy of secondary transplantation sort.**

**A) pooled bone marrow of primary hosts, pre-enriched.**

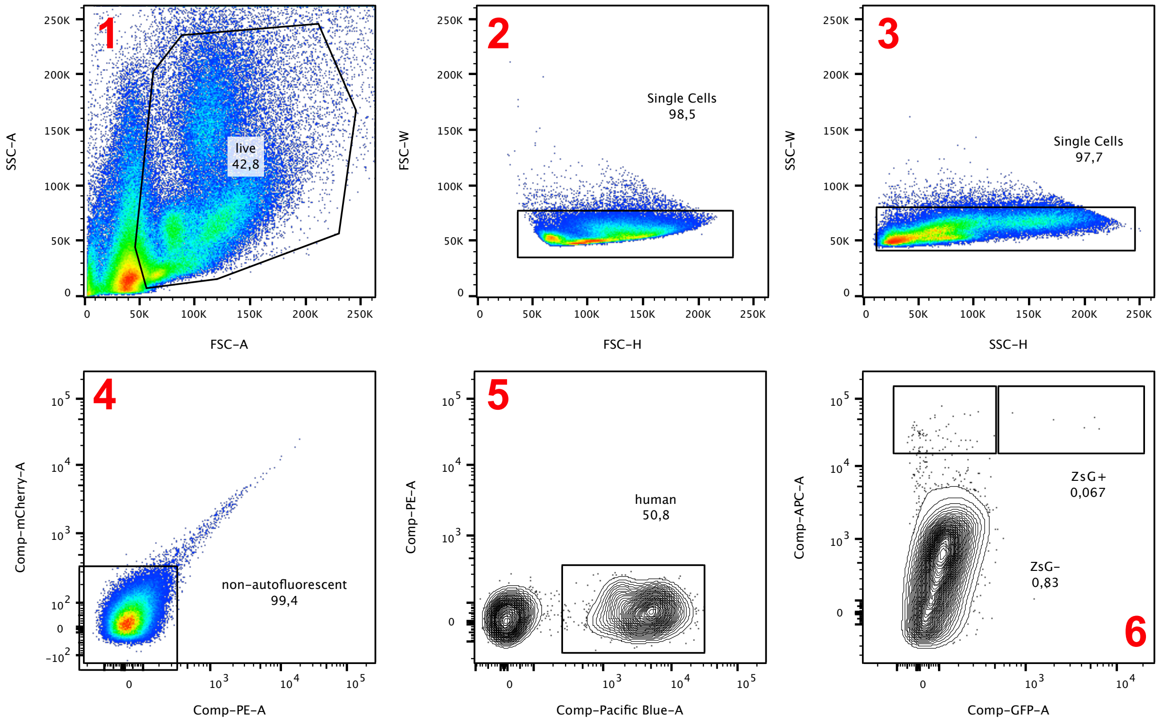


**B) post CD34+ magnetic enrichment.**

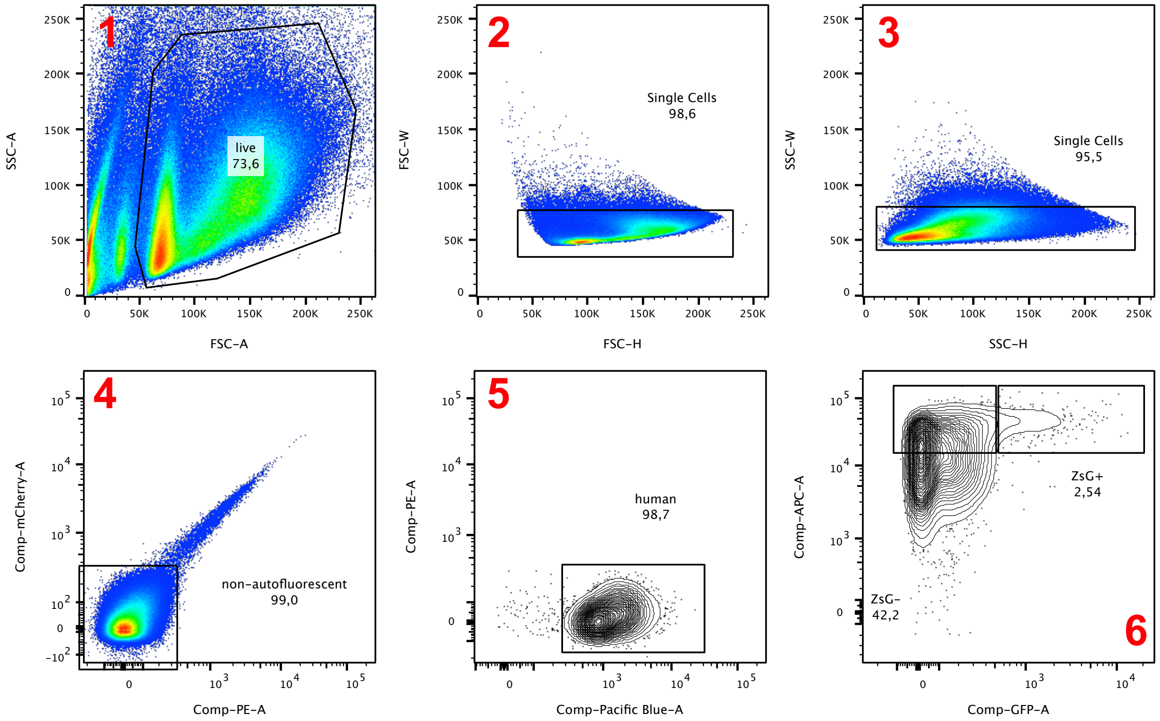
